# Distinct neurophysiological features and memory representations along the long axis of the developing medial temporal lobe

**DOI:** 10.1101/2025.10.14.682334

**Authors:** Qin Yin, Adam J. O. Dede, Robert T. Knight, Eishi Asano, Elizabeth L. Johnson, Noa Ofen

## Abstract

The medial temporal lobe (MTL) is crucial for episodic memory, whereby posterior MTL preferentially represents visuospatial information, and anterior MTL is involved in the representation of semantic or conceptual information. The neurophysiological underpinnings of content-preferential organization in the developing MTL are largely unknown. Here we utilized rare electrocorticography (ECoG) recordings from 23 pediatric epilepsy patients who completed a visual scene recognition memory task to systematically examine the neurophysiological underpinnings of memory formation along the MTL long axis. The timing of high-frequency activity (HFA, ∼70-150 Hz) differed between the posterior and anterior MTL, peaking after scene onset in the posterior MTL and around scene category response (indoor/outdoor scene categorization) in the anterior MTL. Further, in the posterior MTL, HFA was predictive of successful memory formation and positively linked to memory performance, highlighting the importance of posterior MTL HFA to memory formation. In contrast, theta frequency in the anterior MTL was linked to memory performance, and theta-HFA phase-amplitude coupling before scene category responses was predictive of successful memory formation, highlighting the importance of anterior MTL theta oscillations to memory formation. Our findings establish distinct neurophysiological features along the posterior-to-anterior axis of the developing MTL that differentially support the representation of perceptual and conceptual information during memory formation.

## INTRODUCTION

Episodic memory, the ability to remember past experiences, is fundamental to our daily life and continues to develop from childhood into adulthood^1–4^. The medial temporal lobe (MTL) is essential for episodic memory formation^5,6^, supporting memory through a content-based manner, with visuospatial information represented in the posterior region and semantic or conceptual information in the anterior region along its long axis^7–9^. This content-specific organization is well-documented in adults but remains largely unknown whether the developing MTL exhibits similar spatial organization and how the development of this organization might contribute to memory development.

Human functional connectivity^10–12^ and animal tract-tracing^13–15^ studies reveal that the posterior MTL is predominantly connected to regions of the dorsal visual pathway, while the anterior MTL is predominantly connected to regions of the ventral visual pathway. These differential connections may constrain how the MTL preferentially represents visuospatial information, such as scenes, in the posterior region, and stimulus entity, such as objects and faces, in the anterior region^7,8^. Neuropsychological studies provide near-consistent observations of preferential content processing along the MTL long axis, with lesions in the posterior MTL leading to impairments in scene recognition and lesions in the anterior MTL resulting in deficits in recognition for items such as objects and faces^9,16–21^. Neuroimaging studies, however, have yielded mixed findings. While the posterior MTL consistently shows increased responses to successfully remembered scenes^9,22,23^, the anterior MTL exhibits memory-related responses to both items (i.e., objects, faces) and scenes^24–26^. Responses to scenes in the anterior MTL align with its posited role in integrating distinct visuospatial features into higher-level representations such as conceptual categories of scenes^8,15,27–29^. Information feedforward from the posterior MTL to the anterior MTL may underlie the association of visuospatial information in the anterior MTL for the representations of the conceptual aspect of the visuospatial stimuli^13,30^.

Systematic examination of activity along the MTL long axis may enhance our understanding of MTL development. For example, some studies have reported age-invariant MTL involvement for memory for visual scenes^31^ or words^32^, while others have observed age-related increases in posterior MTL involvement for visual scenes^33,34^ or age-related decreases in posterior MTL involvement for objects^35^. The mixed age-related differences and the well-studied content-specific organization along the MTL long axis in adults raise key questions regarding whether the developing MTL exhibits distinct content representations along its long axis. These questions are tightly linked to investigating the underlying neurophysiology along the MTL long axis in the developing brain.

We utilized rare subdural electrocorticography (ECoG) recordings from pediatric epilepsy patients during the performance of an established scene subsequent memory task^31,33,36–42^ to examine the representations of indoor and outdoor visual scenes along the long axis of the developing MTL during memory formation. We hypothesized that the posterior MTL supports visuospatial representation of visual scenes, while the anterior MTL supports conceptual category representation (i.e., whether a stimulus depicts an “indoor” or “outdoor” scene). We used ECoG to define the temporal dynamics and neurophysiological mechanisms associated with external scene onset and internally generated scene category responses. We examined two distinct neurophysiological features: high-frequency activity (HFA, ∼70 – 150 Hz) and oscillations in the theta band (∼2-8 Hz). HFA indexes local neuronal activity^43–47^ and positively correlates with the fMRI BOLD signal^43,45,48–50^, providing a measure of localized neuronal activity with high temporal resolution. Theta is the predominant oscillation in the MTL and has been linked to episodic memory and other MTL-dependent cognitive functions including working memory and spatial navigation^51–54^. It is posited to support MTL computations involved in information integration via the coordination of local activity^55–58^. In addition to supporting local computation, theta also coordinates inter-regional communication and information transfer^59–62^. We examined content representations in the posterior and anterior MTL supporting memory formation by quantifying HFA and theta oscillations associated with scene onset and scene category response onset.

We demonstrate that the posterior and anterior MTL are differentially involved in external scene processing and internal scene category responses, as indicated by the different timing of HFA in each region. The neurophysiological features associated with memory differed between these two regions. The posterior MTL exhibited an HFA-dependent mechanism; increased HFA predicted successful memory formation and was positively linked to memory performance. In contrast, the anterior MTL exhibited a theta-dependent mechanism, where theta frequency correlated with memory performance. Furthermore, in the anterior MTL, increased theta-HFA phase-amplitude coupling before scene category response predicted successful memory formation. Lastly, increased posterior-anterior theta coupling predicted successful memory formation. Our findings establish distinct content representations along the long axis of the developing MTL, supported by different neurophysiological features, which, together, may underlie multifaceted representations of episodic memory.

## RESULTS

### Typical memory performance in pediatric epilepsy subjects

We analyzed ECoG data from 23 subjects (12 females and 11 males; 5.9-20.5 years of age; 13.4 ± 4.0 [M ± SD] years; Table S1) who had seizure-and artifact-free electrodes sampling MTL while they were performing a scene subsequent memory task that has been widely used for studying the neural correlates of memory and memory development^31,33,36–42^. Subjects studied sets of 40 pictures of scenes and responded whether each scene was “indoor” or “outdoor” in preparation for a memory test (Fig. 1a). They were then tested with the previously presented pictures intermixed with 20 new pictures and responded whether they saw the pictures before or not by responding “old” or “new” to each scene. During the study block, a mean of 0.94 ± 0.04 of scenes was correctly identified as indoor/outdoor, indicating that subjects were proficient at correctly categorizing the scenes. During the test block, 0.67 ± 0.17 of the studied scenes were later recognized as “old” (i.e., hit rate), and 0.30 ± 0.17 of the new scenes were later recognized as “old” (i.e., false alarm rate). Memory performance was assessed with recognition accuracy, calculated as the hit rate minus the false alarm rate. Recognition accuracy varied across subjects from 0.02 to 0.76, with a mean ± SD of 0.37 ± 0.21 (Table S1).

Recognition accuracy did not correlate with age in this subsample of patients with MTL coverage (r = 0.20, p = 0.36; Fig. 1b). However, recognition accuracy demonstrated a pattern of age-related increase, reaching significance in data from the larger sample of epilepsy patients we collected regardless of electrode coverage (n = 107; r = 0.26, p = 9.8 × 10^-3^; Fig. 1b). Importantly, we observed large variability in recognition accuracy among adolescents aged ∼12-18 years old, which is comparable to the typical developmental trajectory in large non-clinical samples^31,38,39^ in a similar scene subsequent memory task (n = 186; r = 0.58, p = 3.91 × 10^-18^; Fig. 1b). Altogether, the behavioral pattern suggests typical memory development in our clinical sample, allowing us to investigate neurophysiological signatures supporting memory formation in the developing brain^63^.

**Figure 1.**
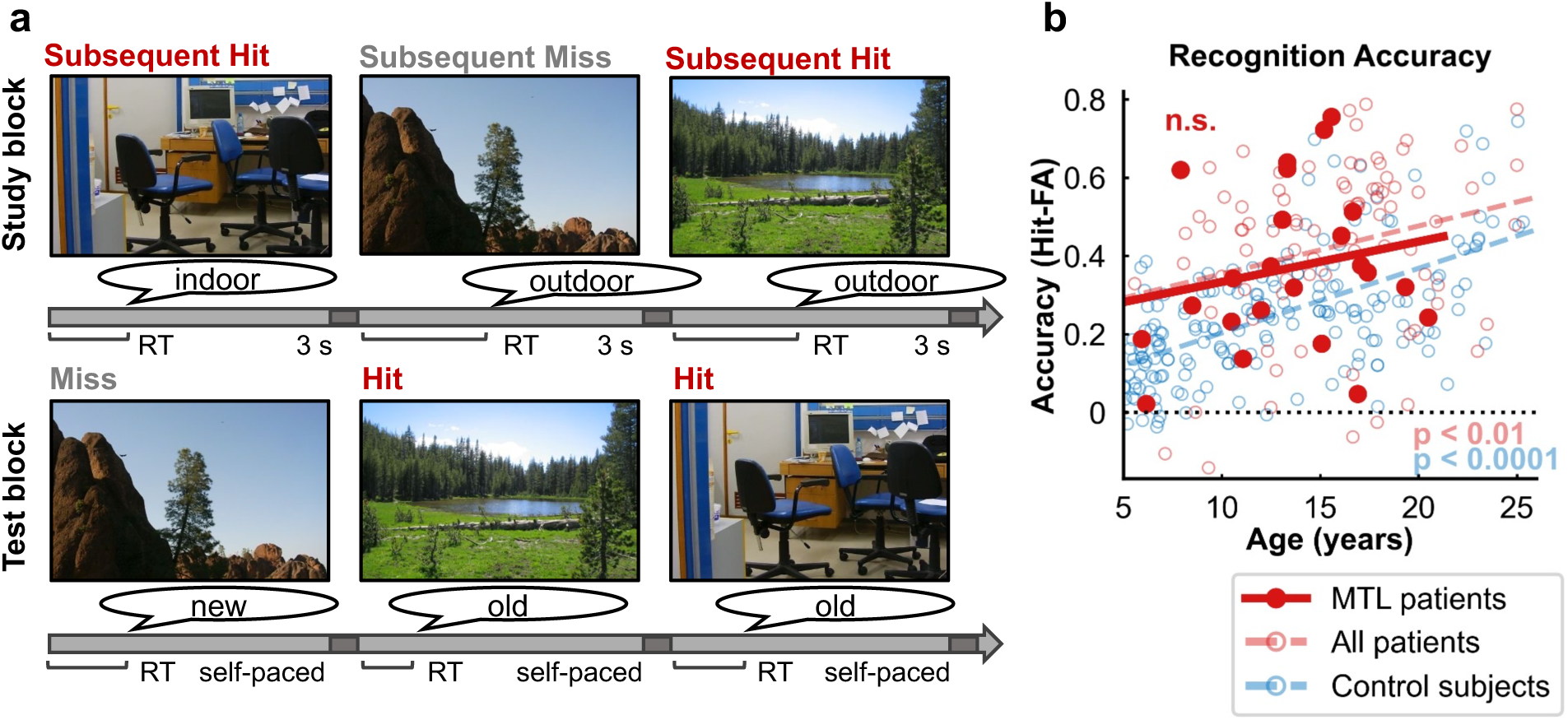
Scene subsequent memory paradigm and memory performance. (**a**) Subsequent memory paradigm. Pictures of scenes were presented in two consecutive study-test cycles. In each study cycle, subjects studied 40 scenes (3 s each, separated by a 0.5-s interstimulus fixation) and indicated whether each scene was indoor/outdoor. In each test cycle, subjects were tested on all 40 studied scenes, intermixed with 20 new scenes as foils. Subjects made an old/new judgment of each scene based on whether they remembered seeing the scene at study. The test was self-paced. Each response at the test was coded as a hit (old response to a studied scene), miss (new response to a studied scene), correct rejection (new response to a foil scene), or false alarm (old response to a foil scene). Based on the test response, scenes at the study cycle were categorized as subsequent hit or subsequent miss. (**b**) Recognition accuracy by age. Epilepsy patients with MTL electrode coverage (red filled circle and solid line) demonstrate a similar pattern of age-related increase in recognition accuracy (hit rate – false alarm rate) as all epilepsy patients (red empty circle and dashed line) and normal developing control subjects (blue empty circle and dashed line).

### HFA peak latency increases along the MTL posterior-to-anterior axis

HFA has been used to investigate the temporal dynamics of brain activity during cognitive tasks in adults^64–68^ and the developing brain^37^. We analyzed ECoG signals from 90 artifact-free and nonpathological surface MTL electrodes [median(IQR), 4(2) electrodes/subject] across 23 subjects. A time-frequency representation of power for each study trial was computed from 70 to 150 Hz across 30 logarithmically spaced frequency bands, z-scored on the pre-stimulus baseline via statistical bootstrapping^37,68^ for each frequency band, and then averaged across frequencies. Trials with incorrect indoor/outdoor responses were excluded, partly controlling for the effects of attention during study on memory formation.

We plotted HFA across all MTL electrodes ordered from posterior to anterior (Fig. 2a) and observed that scene viewing induced increased HFA in both subsequent hit and subsequent miss trials (Fig. 2a; z > 2.75, p < 0.01). Moreover, the temporal dynamics of HFA differed between posterior and anterior electrodes. In posterior electrodes, HFA increased after scene onset and persisted throughout the first second of scene viewing (Fig. 2a). In anterior electrodes, HFA increased later during scene presentation (Fig. 2a). Spatiotemporal dynamics along the posterior-to-anterior axis were quantified with the correlation between HFA peak latencies and the y-coordinates of electrodes. Here, HFA peak latency was defined as the time of maximum HFA between scene onset and 0.2 s after scene category response onset on each trial (see Fig. 2b for examples) and averaged across trials. A linear mixed-effects model with subsequent memory (i.e., hit vs. miss) and y-coordinate as fixed effects, and subjects and nested electrodes as random effects revealed a main effect of y-coordinate (β = 0.66, t(176) = 3.24, p = 1.4 × 10^-3^) and subsequent memory × y-coordinate interaction (β = -0.31, t(176) = -2.45, p = 0.02). No main subsequent memory effect was observed (β = 0.10, t(176) = 0.76, p = 0.44). Post-hoc analysis using separate linear mixed-effect models for subsequent hit and miss trials, with y-coordinate as the fixed effect and subjects and nested electrodes as random effects, revealed that HFA peak latency increased from posterior to anterior along the MTL long axis in subsequent hit (Fig. 2c, left; β = 0.31, t(88) = 4.44, p = 2.59 × 10^-5^) but not in subsequent miss trials (Fig. 2c, right; β = 0.07, t(88) = 0.61, p = 0.55).

The different posterior and anterior HFA patterns (Fig. 2a) and increased HFA peak latency from posterior to anterior (Fig. 2b) support different processes along the MTL long axis. We thus continued to investigate functional differentiation along the MTL long axis in the developing brain by defining posterior and anterior MTL regions based on the localization of MTL electrodes across subjects. We classified MTL electrodes as posterior or anterior using the midpoint between the most anterior and most posterior electrodes (Fig. 2d, y = -14.35 mm). This approach allowed utilizing the extant data and avoids potential biases in defining functional subregions along the MTL long axis^22,23,26^. Posterior MTL activity was sampled from 29 electrodes [median(IQR), 1(1) electrodes/subject] across 17 subjects (13.25 ± 3.68 years), and anterior MTL activity was sampled from 61 electrodes [median(IQR), 3(2) electrodes/subject] across 20 subjects (12.98 ± 3.88 years).

**Figure 2.**
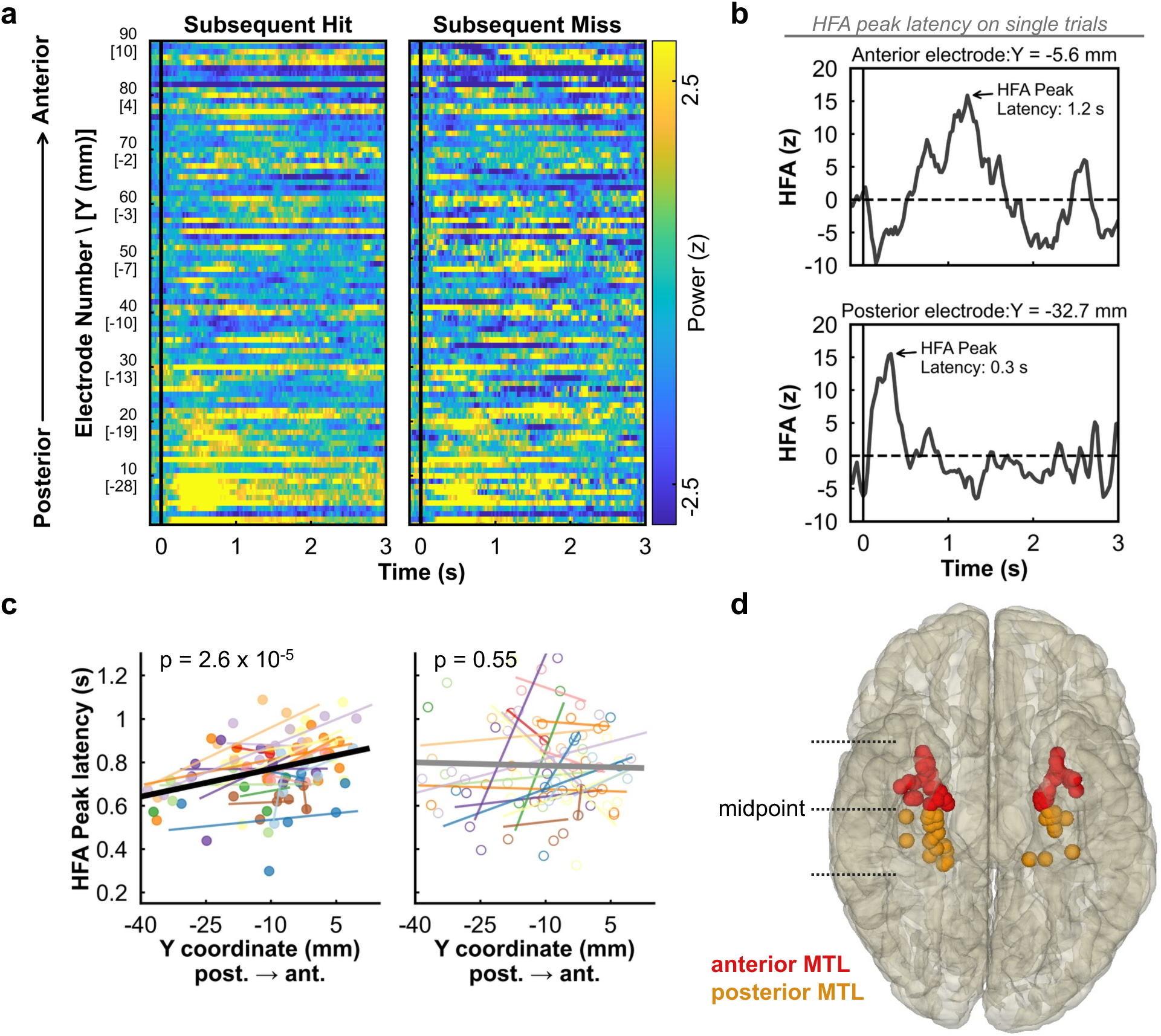
Spatiotemporal dynamics of HFA along the MTL long axis. (**a**) HFA across all MTL electrodes sorted by y-coordinates, ordered from posterior to anterior. The solid vertical line indicates scene onset. Scene viewing induced increased HFA (z > 2.57, p < 0.01) in the MTL in both subsequent hit (left) and miss trials (right). The HFA increases occurred in the first second of scene viewing in posterior electrodes but were present later during the scene presentation in anterior electrodes. (**b**) Example trials showing HFA peak latency definition from an anterior (top) and a posterior (bottom) electrode. (**c**) HFA peak latency increased along the MTL long axis in subsequent hit trials (left) but not in subsequent miss trials (right). Each color represents a single subject. (**d**) Electrode coverage of the posterior and anterior MTL from all subjects. Electrodes were classified as posterior or anterior using the midpoint between the most anterior and most posterior electrodes. Orange, posterior MTL; red, anterior MTL.

### HFA in the anterior MTL tracks scene category response

We next examined the temporal dynamics of HFA relative to scene onset and scene category responses (i.e., indoor/outdoor) in the posterior and anterior MTL. We first visualized HFA on the single-trial level by averaging across electrodes within each region per subject, pooling trials across all subjects, and sorting them by response time (RT). We observed that HFA in the posterior MTL increased after stimulus onset and was sustained throughout the first second of scene viewing regardless of RT in subsequent hit trials (Fig. 3a). In contrast, HFA in the anterior MTL did not increase following stimulus onset but instead increased around each trial’s RT in both subsequent hit and miss trials (Fig. 3b). HFA peak latency was again used to assess the relationship between HFA and RT. A linear mixed-effects model with region (posterior MTL vs. anterior MTL), subsequent memory (hit vs. miss), and RT as fixed effects, and subjects and nested electrodes as random effects, revealed a significant region × RT interaction (β = -0.41, t(172) = -2.08, p = 0.04) and region × RT × subsequent memory interaction (β = 0.27, t(172) = 2.22, p = 0.03). There was no significant region, subsequent memory, RT, region × subsequent memory, or subsequent memory × RT effects (Table S2).

Post hoc analyses were conducted with per-region linear mixed-effects models with subsequent memory and RT as fixed effects. In the posterior MTL, there were no significant subsequent memory or RT effects on HFA peak latency (Table S3). However, we observed a significant subsequent memory × RT interaction (Fig. 3c; β = 0.48, t(54) = 3.32, p = 1.6 × 10^-3^), with HFA peak latency tracking RT in subsequent miss (β = 0.69, t(27) = 4.85, p =4.5 × 10^-5^) but not subsequent hit (β =0.21, t(27) = 1.74, p = 0.09) trials, suggesting scene onset-induced activity rather than response-related activity in the posterior MTL is beneficial for successful memory formation. In the anterior MTL, we observed a significant RT effect (Fig. 3d; β = 0.74, t(118) = 3.04, p = 2.9 × 10^-3^) in that the anterior MTL HFA tracked subjects’ scene category responses. We did not observe significant subsequent memory or subsequent memory × RT interaction effects (Table S3). These findings suggest an endogenous process in the anterior MTL, likely reflecting conceptual category representation of visual scenes.

**Figure 3.**
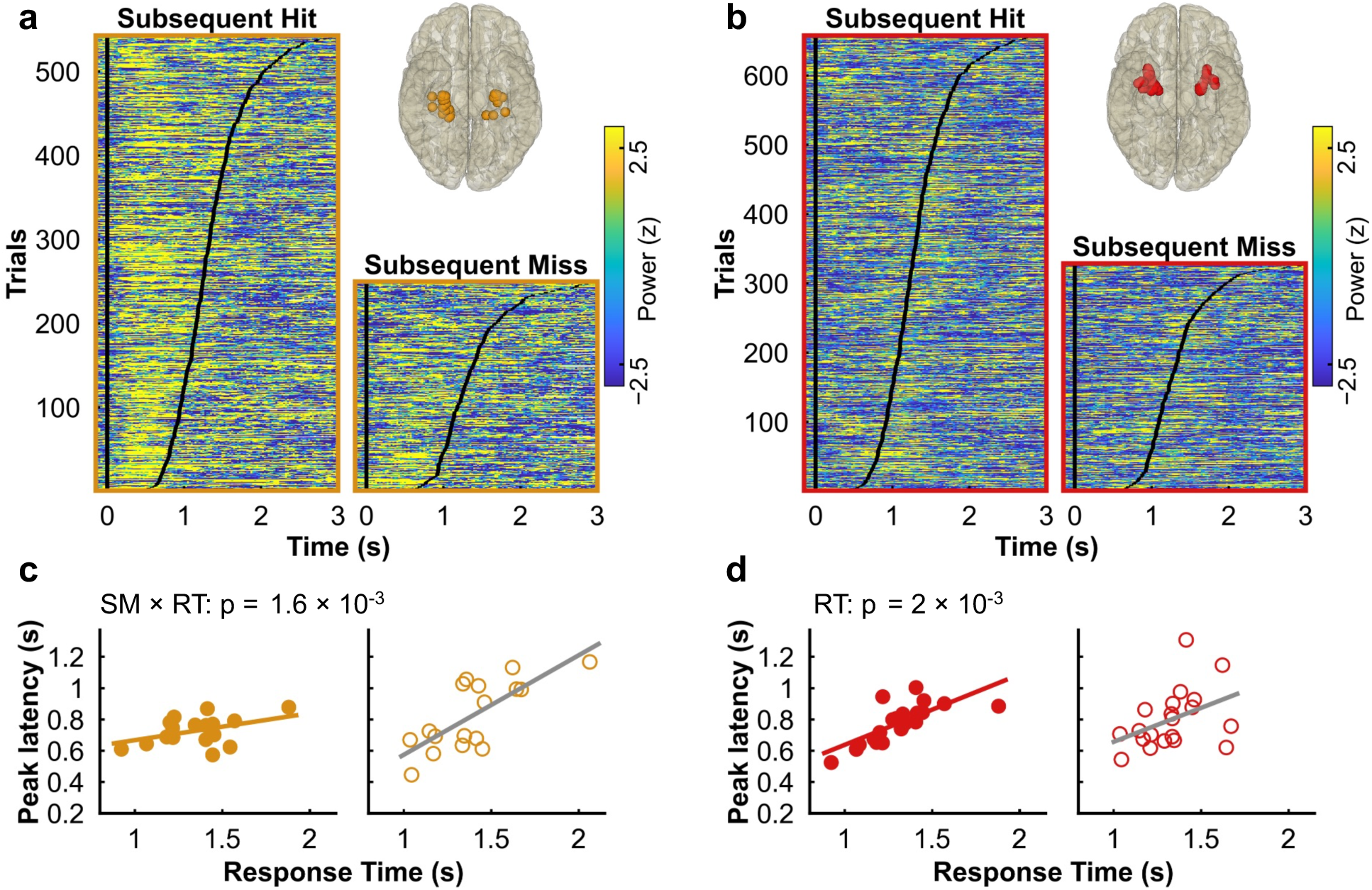
Associations of HFA peak latency with scene category RT. (**a**) HFA across all trials in the posterior MTL, sorted by RT (black tick marks) from the fastest to slowest response trial in subsequent hit (left) and miss trials (right). Each trial represents data averaged across electrodes within the posterior MTL per subject. The solid vertical line indicates the scene onset. (**b**) Same as (a) in the anterior MTL. (**c**) HFA peak latency in the posterior MTL was not associated with RT in hit trials (left) but showed a positive association in subsequent miss trials (right). P-value indicates subsequent memory × RT interaction. (**d**) HFA peak latency in the anterior MTL positively associated with RT in both subsequent hit (left) and miss (right) trials. P-value indicates the main effect of RT.

### HFA in the posterior MTL predicts memory formation and is associated with recognition accuracy

After identifying different temporal dynamics of activity in the posterior and anterior MTL, we further investigated the subsequent memory effects (SMEs) by contrasting the HFA of subsequent hit trials with subsequent miss trials. We analyzed SMEs by aligning the trials to scene onset for scene-induced SMEs and by aligning the trials to subjects’ scene category response onset for response-related SMEs. SMEs were analyzed between -0.15 to 1 s for stimulus-locked data and between -1 to 0.2 s for response-locked data. We conducted each subsequent memory analysis with a linear mixed-effects model with subsequent memory (hit vs. miss) as the fixed effect and subjects and nested electrodes as random effects and used cluster-based correction for multiple comparisons. In the posterior MTL, we observed increased HFA during successful memory formation in both stimulus-locked data (i.e., positive SMEs, 0.475 to 0.625 s, p = 0.03; Fig. 4a, top) and response-locked data (-0.475 to -0.375 s, p = 0.04, Fig. 4a, bottom). These findings provide evidence that posterior MTL HFA is beneficial for successful memory formation of visual scenes. In the anterior MTL, we observed clusters of positive SMEs in both stimulus-locked data (0.650 to 0.775 s, p = 0.02; Fig. 4b, top) and response-locked data (-0.425 to -0.275 s, p = 8 × 10^-3^; Fig. 4b, bottom). Of note, these positive SMEs in the anterior MTL were driven by decreased HFA in subsequent miss trials.

We then examined whether the observed SMEs were associated with memory performance or age. We averaged across time points within each cluster of SMEs per subject and submitted the averaged data into multiple regression with recognition accuracy and age as predictors. In the posterior MTL, scene-induced SME was not associated with recognition accuracy or age (Fig. 4c, top; Table S4). However, response-locked SME was positively associated with recognition accuracy (Fig. 4c, bottom; β = 0.56, t(13) = 2.57, p = 0.02). We observed no age or age × recognition accuracy interaction effects for response-locked SME (Table S4), demonstrating that HFA in the posterior MTL supports memory independent of age. In the anterior MTL, we did not observe recognition accuracy, age, or age × recognition accuracy interaction effects on either scene-induced or response-related SMEs (Fig. 4d; Table S4). Taken together, our findings demonstrate that the posterior MTL supports scene processing across age, with HFA serving as a neurophysiological feature linked to memory formation of visual scenes.

**Figure 4.**
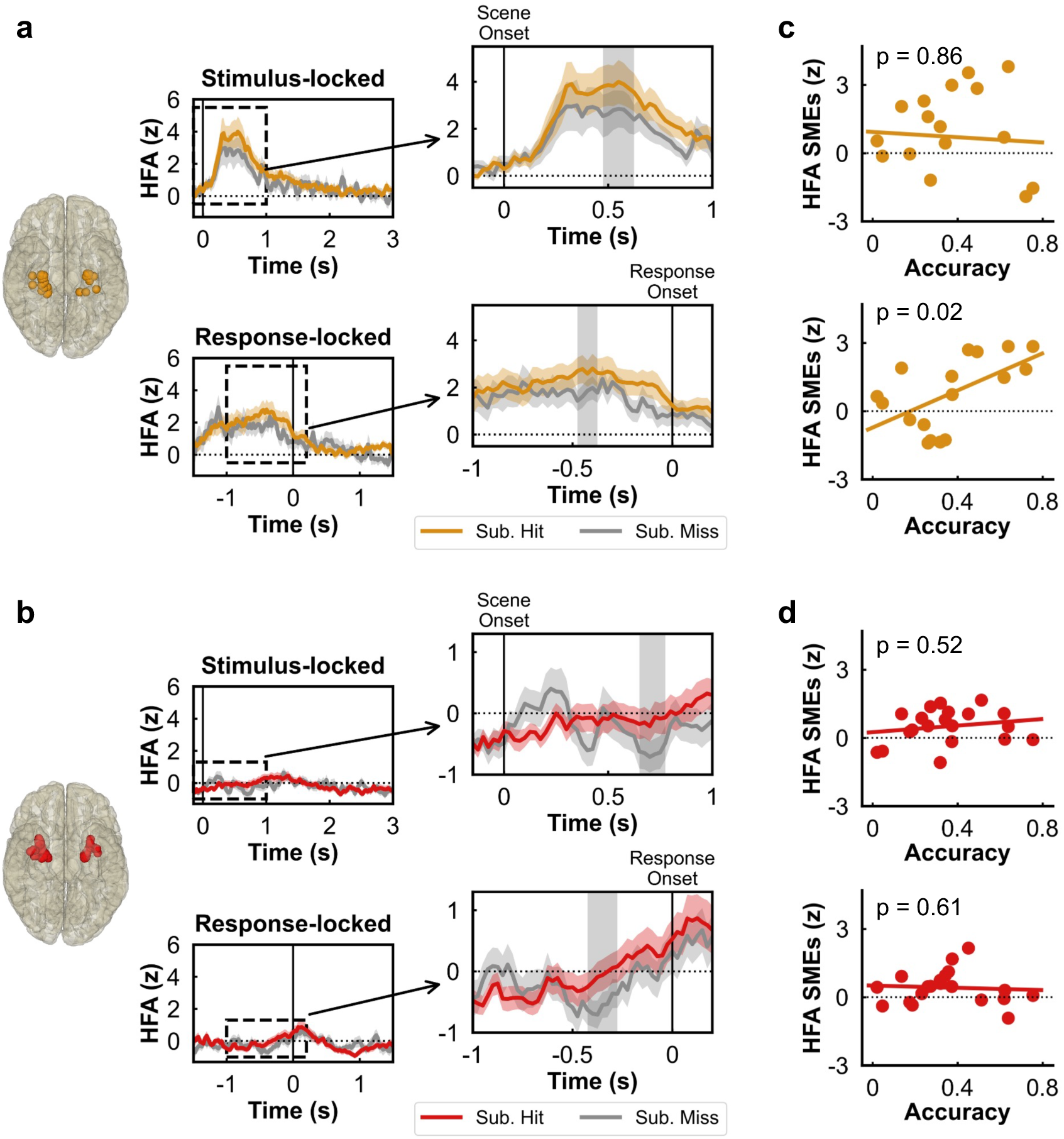
HFA signatures of successful memory formation. (**a**) HFA SMEs in the posterior MTL. Top: HFA locked to scene onset. The solid vertical line indicates the scene onset. The inset on the right zooms in on the first second after scene onset, indicating the time window for which SMEs were analyzed. Bottom: HFA locked to scene category response onset. The solid vertical line indicates response onset. The inset on the right zooms in on the one second before the response, indicating the time window for which SMEs were analyzed. HFA data were averaged across subjects, and the shaded areas represent the standard error of the mean. The gray shaded area in the insets denotes cluster-corrected significant SMEs. Sub. Hit, subsequent hit; Sub. Miss, subsequent miss. (**b**) Same as (a) in the anterior MTL. (**c**) Top: scene-induced SME in the posterior MTL was not associated with recognition accuracy. Bottom: response-related SME in the posterior MTL was positively associated with recognition accuracy. The SMEs are grey shaded area in (a). (**d**) Scene-induced (top) and response-related (bottom) SMEs were not associated with recognition accuracy. The SMEs are grey-shaded areas in (b).

### Theta oscillations in the anterior MTL are associated with recognition accuracy

Theta is the predominant oscillation in the MTL and is posited to support MTL computations involved in information integration^55–57^. Emerging evidence suggests that neural oscillations, including theta, show variability in center frequency and power across brain regions^69–71^, task states^72,73^, and development^36,40,42,63,74–76^ and aging^77–79^. To characterize theta variability across MTL subregions and individuals, we utilized a widely used oscillation detection algorithm, SpecParam^80^, to isolate and parameterize oscillations (see Fig. 5a for an electrode example from the anterior MTL). Neural oscillations are defined as peaks above the 1/f aperiodic component in the power spectrum (Fig. 5a, top), and each peak can be parameterized with measures of center frequency, power, and bandwidth (Fig. 5a, bottom). Oscillation detection was conducted between 1-30 Hz per electrode on the average power spectrum, separately for subsequent hit and miss trials. As shown in Fig. 5b, electrodes in the posterior and anterior MTL exhibited oscillatory peaks in both subsequent hit and miss trials. The frequency distribution of the largest peak, i.e., peak with the largest power, across all electrodes per region was examined to identify the frequency range of the dominant oscillation (Fig. 5c). We observed that the largest peaks clustered between ∼1-10 Hz in both the posterior [Fig. 5c; median(IQR): subsequent hit, 7.59(7.67) Hz; subsequent miss, 7.61(2.23) Hz] and anterior MTL [Fig. 5c; median(IQR): subsequent hit, 6.93(6.14) Hz; subsequent miss: 7.11(6.78) Hz], confirming that theta oscillations dominate the developing MTL^36^. We selected the largest peak between 1-10 Hz as theta oscillation per electrode based on the frequency distribution for further analysis.

We examined whether theta parameters (i.e., frequency, power, and bandwidth, see Fig. 5a) differed between the posterior and anterior MTL and between subsequent hit and miss trials. Linear mixed-effects models were used per parameter with region (posterior MTL vs. anterior MTL) and subsequent memory (hit vs. miss) as fixed effects and subjects and nested electrodes as random effects. Theta frequency did not differ between regions or between subsequent hit and miss trials (see Fig. S1; Table S5). However, we observed a significant negative SME on theta power (Fig. 5d; β = -0.11, t(175) = -2.95, p = 3.7 × 10^-3^), extending previous findings of broadband power decreases during successful memory formation^36,42^ to narrowband theta oscillations. No significant region or region × subsequent memory effect was observed on theta power (Fig. 5d; Table S5). Theta bandwidth also did not differ between regions or between subsequent hit and miss trials (Fig. S1; Table S5). The lack of regional differences across parameters is consistent with the idea that a common theta frequency may provide a functional basis for the interaction and information transformation between these two regions^71,81^.

We then examined individual differences. Each theta parameter was submitted to a linear mixed-effects model per region with age, recognition accuracy, and subsequent memory as fixed effects, and subjects and nested electrodes as random effects. In the posterior MTL, no significant effects were observed on theta frequency (see Fig. S2a; Table S6). However, we observed significant effects of age and subsequent memory on theta power (Fig. 5e; age: β = 0.47, t(50) = 2.15, p = 0.04; subsequent memory: β = - 0.12, t(50) = -3.12, p = 3.0 × 10^-3^). No other significant effects were observed for theta power (see Fig. S2a; Table S6), and no significant effects were observed for theta bandwidth (see Fig. S2a; Table S6). Taken together, we found that, in the posterior MTL, theta power increased with age, possibly reflecting the reduced involvement of theta with development.

In the anterior MTL, we observed that higher theta frequency was associated with better memory performance (Fig. 5f; β = 0.42, t(113) = 2.77, p = 6.6 × 10^-3^). No other significant effects on theta frequency were observed (see Fig. S2b; Table S7). For power, we observed a significant SME (β = -0.11, t(113) = -3.00, p = 3.3 × 10^-3^). No other significant effects were observed for power (see Fig. S2b; Table S7), and no significant effects were observed for theta bandwidth (see Fig. S2b; Table S7). Taken together, we found that, in the anterior MTL, faster theta oscillations were associated with better memory performance independent of age, suggesting increased precision of activity may support better memory in the developing brain^42^.

**Figure 5.**
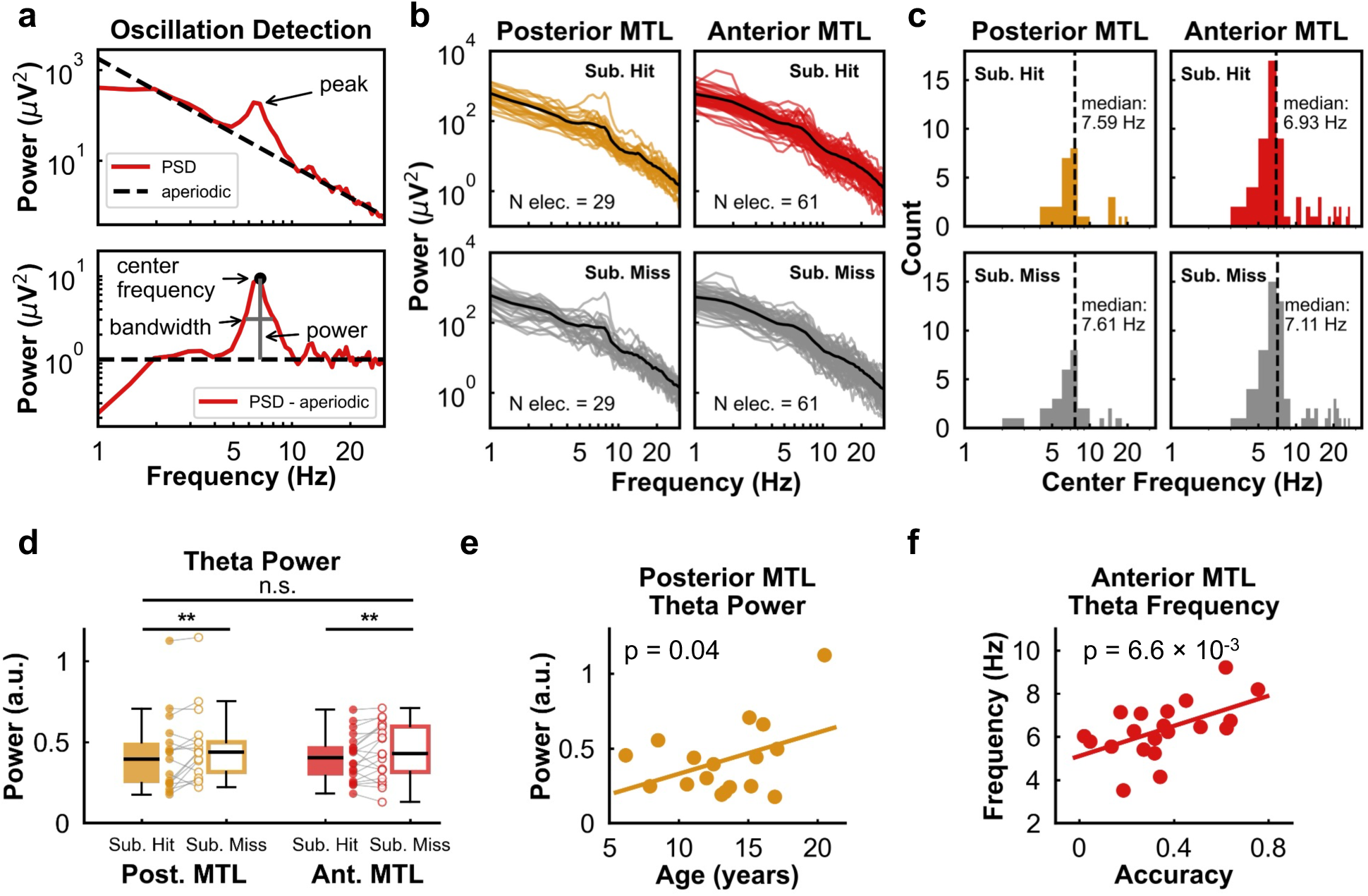
Characterizing theta oscillations in the posterior and anterior MTL. (**a**) Top: power spectrum of one representative electrode from the anterior MTL showing the existence of an oscillation as a peak over the estimated aperiodic activity (black dashed line). Bottom: power spectrum after subtracting the estimated aperiodic activity. The detected oscillation is parametrized with center frequency, power, and bandwidth. (**b**) Power spectra across all electrodes for subsequent hit (top) and miss (bottom) trials. The black line in each plot indicates the averaged power spectrum across all electrodes within each region. Sub. Hit, subsequent hit; Sub. Miss, subsequent miss. (**c**) Center frequency distribution of the largest peak between 1-30 Hz shows that theta dominates the MTL. (**d**) Theta power between regions and between subsequent hit and miss trials. ** p ≤ 0.01. Post. MTL, posterior MTL; Ant. MTL, anterior MTL. (**e**) Posterior MTL theta power increased with age. (**f**) Anterior MTL theta frequency was positively associated with recognition accuracy.

### Anterior MTL theta-HFA coupling before scene category response predicts memory formation

The association of theta oscillations with memory performance suggests that theta-related mechanisms underlie memory. We next examined theta-HFA phase-amplitude coupling, a mechanism that has been posited to support information binding and mnemonic representations^55–58^. Examining theta-filtered and HFA-filtered signals on the single-trial level (see Figs. 6a, 6b for examples), we observed theta phase-dependent variations in HFA, and the dependence varied across time. To capture the temporal dynamics of theta-HFA coupling, we analyzed event-related theta-HFA PAC (ERPAC). ERPAC was calculated with theta defined by the individually detected frequency per electrode for subsequent hit and miss trials. This approach allows the capture of coupling strength while controlling the inter-individual differences in theta parameters^36,42,80^. Building on the distinct temporal dynamics of the posterior and anterior MTL revealed by HFA, we again investigated SMEs by aligning the studied trials to scene onset for scene-induced effects and scene category response onset for response-related effects. Subsequent memory analysis was conducted using linear mixed-effects model with cluster-based correction. We did not observe any SMEs in the posterior MTL in either stimulus-locked data or response-locked data (Fig. 6c). In the anterior MTL, no SMEs were observed in stimulus-locked data (Fig. 6d, top). However, a cluster of positive SMEs (-0.470 to -0.354 s, p = 0.04; Fig. 6d, bottom) was observed in response-locked data. These positive SMEs suggest that theta-HFA coupling in the anterior MTL, occurring before subjects’ scene category responses, underlies conceptual scene category representation and facilitates successful memory formation.

After identifying positive SMEs in the anterior MTL, we examined whether this signature of memory formation was associated with age or recognition accuracy. We averaged across time points within this cluster of positive SMEs (Fig. 6d, bottom) per subject and submitted the averaged ERPAC into multiple regression with recognition accuracy and age as predictors. We did not observe any significant effects of age or recognition accuracy (Table S8). Controlling for individual differences in theta parameters, the lack of individual differences in ERPAC indicates that theta frequency (see Fig. 5f), but not the strength of theta-HFA coupling, explains individual differences in recognition accuracy, suggesting that increased precision of the coupling mechanism supports better memory in the developing brain^42^.

Lastly, theta supports memory by operating through distinct theta phases^82–84^. To capture the theta phase associated with increased HFA in the anterior MTL, we analyzed the preferred phase of HFA during the time window of significant SMEs (see Fig. 6d, bottom). Theta phases were divided into 18 bins from −π to π, and HFA was binned based on the theta phases. We observed increased HFA at the theta trough during subsequent hit trials (Figs. 6e, 6f). In contrast, during subsequent miss trials, HFA increased at both the trough and midpoint. Together, these observations suggest that theta trough at the anterior MTL provides the optimal cyclical window for neuronal activity and underlies successful memory formation.

**Figure 6.**
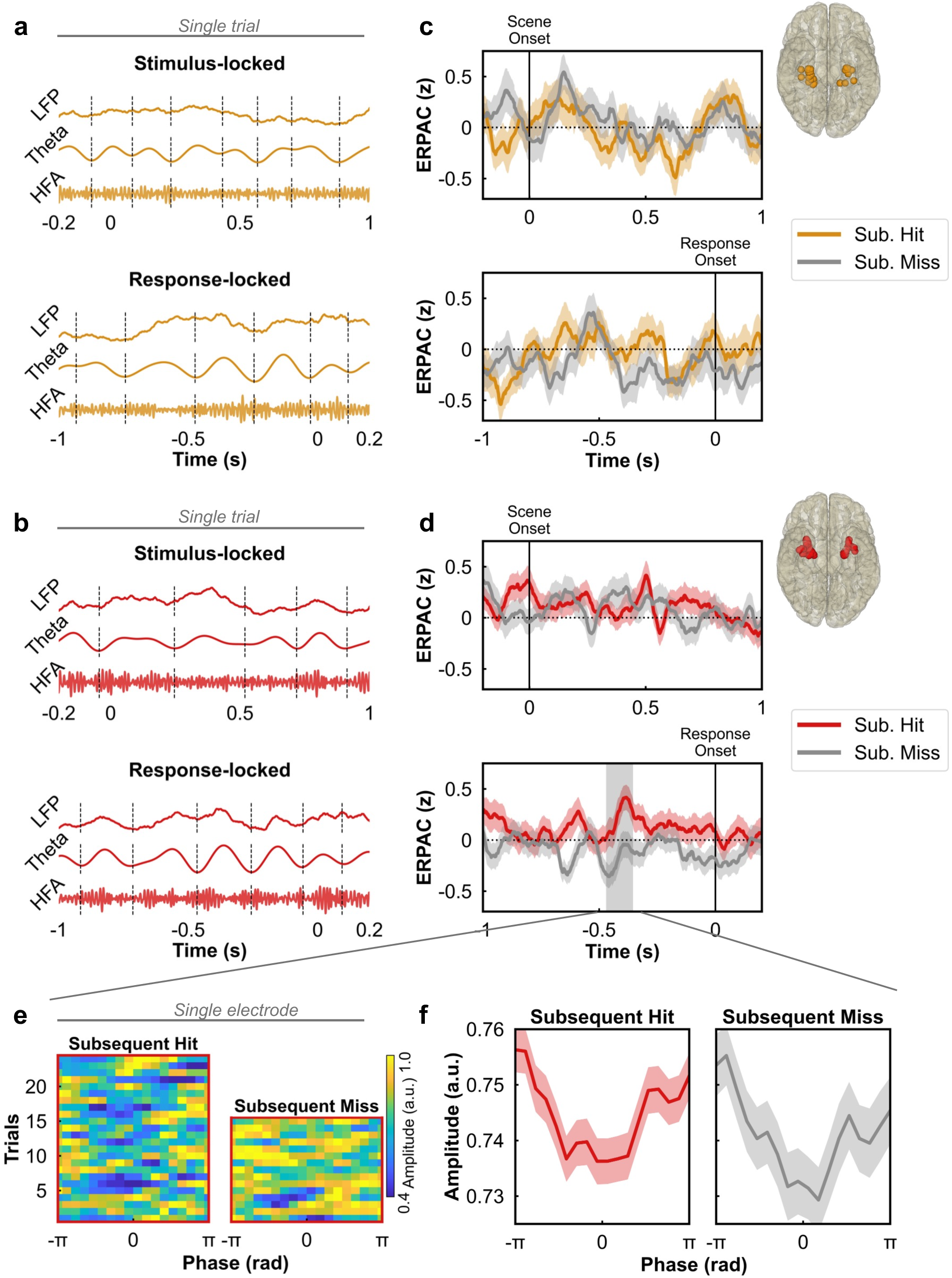
Anterior MTL theta-HFA coupling associated with successful memory formation. (**a**) Raw, theta-filtered, and HFA-filtered signals of a representative subsequent hit trial from a posterior MTL electrode for stimulus-locked data (top) and response-locked data (bottom). Dashed vertical lines indicate theta troughs. (**b**) Same as (a) from an anterior MTL electrode. (**c**) ERPAC for subsequent hit and miss trials across all subjects in the posterior MTL. Top: ERPAC locked to stimulus onset. The solid line at time zero indicates stimulus onset. Bottom: ERPAC locked to response onset. The solid line at time zero indicates “indoor/outdoor” response onset. Shaded areas represent the standard error of the mean across subjects. No significant SMEs were observed in the posterior MTL. Sub. Hit, subsequent hit; Sub. Miss, subsequent miss. (**d**) Same as (c) for the anterior MTL. A cluster of positive SMEs was observed preceding response onset (bottom). The gray shaded area denotes cluster-corrected significant SMEs. (**e**) HFA from the time window of significant SMEs (Fig. 6d, bottom) binned by theta phases across subsequent hit (left) and miss trials (right) from a representative anterior MTL electrode. (**f**) HFA from the time window of significant SMEs (Fig. 6d, bottom) binned by theta phases across all subjects for subsequent hit (left) and miss trials (right). Shaded areas indicate the standard error of the mean across subjects.

### Posterior-anterior MTL theta phase coupling predicts memory formation

Having identified distinct neurophysiological features underlying memory formation in the posterior and anterior MTL, we last sought to examine the interaction between these two subregions. Oscillation phase coupling is posited as a mechanism to support inter-regional interaction and information transfer^59,62^. The consistency in theta parameters between the posterior and anterior MTL (See Fig. 5d, Fig. S1) may provide a functional basis for their interaction. Theta phases between the posterior and anterior MTL were aligned during single trials (Fig. 7a), and the phase-locking value (PLV; Fig. 7b) was analyzed to measure theta phase coupling between these two regions. Here, PLV was calculated with the individually detected theta frequency per electrode for subsequent hit and miss trials. SMEs were analyzed using linear mixed-effects models with subsequent memory (hit vs. miss) as the fixed effect and subjects and nested electrodes as random effects. We observed two clusters of positive SMEs in stimulus-locked data (Fig. 7b, top; cluster 1, -0.200 to 0.144 s from scene onset, p = 1 × 10^-3^; cluster 2, 0.271 to 1.000 s from scene onset, p < 1 × 10^-3^) and one cluster of positive SMEs in response-locked data (Fig. 7b, bottom; -0.866 to 0.200 s from scene category response onset, p < 1 × 10^-3^), showing that sustained interaction between the posterior and anterior MTL supports successful memory formation.

We then examined whether any of these SMEs were associated with age or recognition accuracy. The average SMEs per subject for each cluster were submitted to multiple regression with accuracy and age as predictors. We did not observe any significant effects for stimulus-locked or response-related SMEs (Table S9). Controlling for individual differences in theta frequency, these findings indicate that increased precision (see Fig. 5f), rather than the strength, of inter-regional theta coupling is associated with better memory performance^42^.

Lastly, we examined whether posterior MTL HFA were coupled to anterior MTL theta phase at the individually defined frequency. We analyzed inter-regional theta-HFA coupling with ERPAC. Linear mixed-effects models revealed no significant SMEs in either stimulus-or response-locked data (see Fig. S3). However, in the exploratory analysis of examining the correlation between posterior MTL HFA and anterior MTL theta, we observed significant correlations. Posterior MTL HFA during the stimulus-locked SMEs time window (see Fig. 4a, top) was positively correlated with anterior MTL theta frequency in both subsequently hit and miss trials (Fig. 7c, left; hit, r = 0.68, r = 6.8 × 10^-3^; miss, r = 0.65, p = 0.01). During the response-locked SMEs time window (see Fig. 4a, bottom), posterior MTL HFA was positively correlated with anterior MTL theta frequency in subsequent hit trials (Fig. 7c, right; r = 0.60, p = 0.02), but not in miss trials (Fig. 7c, bottom; r = 0.36, p = 0.21). No other correlations between posterior MTL HFA and anterior MTL theta parameters were observed (see Figure S4; Table S10). Taken together, these findings suggest that inter-regional theta coupling facilitates the transfer of information and supports the representation of both the details and the conceptual category of visual scenes during memory formation.

**Figure 7.**
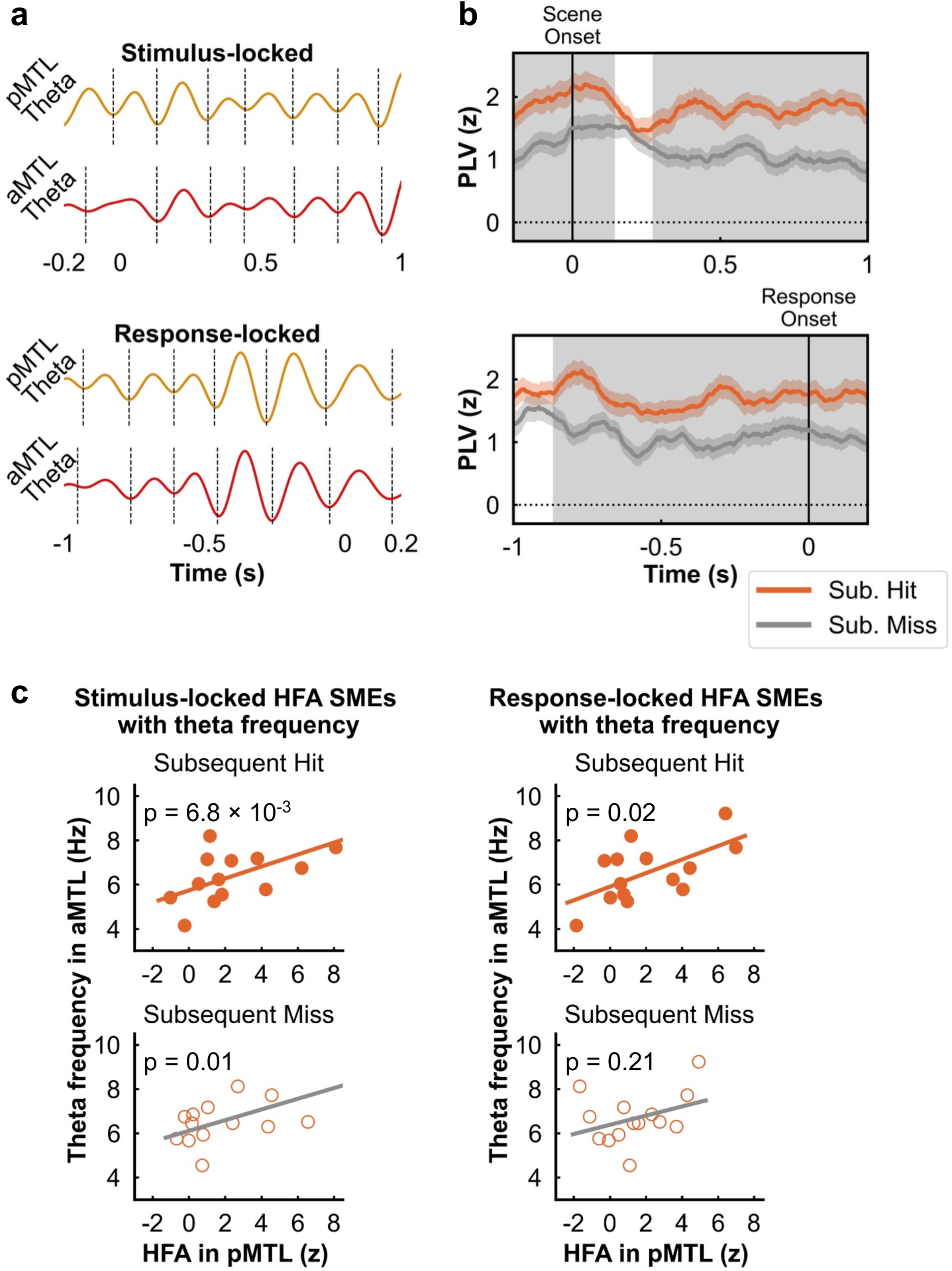
Theta coupling associated with successful memory formation. (**a**) Top: theta-filtered signal of a representative subsequent hit trial from a posterior (top) and anterior (bottom) MTL electrode for stimulus-locked data. Bottom: The same signal locked to the scene category response onset. Dashed vertical lines indicate theta troughs. (**b**) Posterior-anterior MTL theta coupling in stimulus-locked (top) and response-locked (bottom) data. Shaded areas represent the standard error of the mean across subjects. The gray shaded areas denote cluster-corrected significant SMEs. Sub. Hit, subsequent hit; Sub. Miss, subsequent miss. (**c**) Correlation of posterior MTL HFA with anterior theta frequency for subsequent hit (top) and miss (bottom) trials. Left: HFA from the time window of significant SMEs from stimulus-locked data, see Fig. 4a, top, grey shaded area. Right: HFA from the time window of significant SMEs from response-locked data, see Fig. 4a, bottom, grey shaded area.

## DISCUSSION

Human episodic memory is a multifaceted construct that relies on the representations of both perceptual and conceptual information about experiences^85–87^. Utilizing the high spatiotemporal precision afforded by ECoG, we investigated how the developing MTL supports memory formation for visual scenes. We focused our investigation on identifying features that correspond to representations of perceptual details and conceptual category of visual scenes during memory formation, defined by their timing relative to the scene onset and categorization response. Distinct neurophysiological features were observed in the posterior and anterior MTL, suggesting complementary representations of perceptual and conceptual information during memory formation in the developing MTL.

### HFA peak latency indicates different processes in the posterior and anterior MTL

Leveraging the high spatiotemporal resolution of HFA, we observed that HFA peak latency increased along the posterior-to-anterior axis during subsequent hit but not miss trials, suggesting information transfer along the MTL long axis during successful memory formation. Early posterior activity likely reflects responses to visual scenes^9,22,23^, whereas later anterior activity supports the posited role of the anterior region in integrating incoming visuospatial information for higher-level representations^8,15,27–29^. To test whether these dynamics tracked scene onset or scene category responses (i.e., indoor/outdoor), we divided the MTL into posterior and anterior regions. In the posterior MTL, scene onset induced HFA increases at the single-trial level during subsequent hit but not miss trials (Fig. 3a). Moreover, HFA peak latency during subsequent miss but not hit trials correlated with subjects’ RTs. These findings suggest that scene-induced posterior MTL HFA supports successful memory formation.

By contrast, in the anterior MTL, HFA peaked around subjects’ scene category responses at the single-trial level (Fig. 3b), and peak latencies correlated with subjects’ RTs. This category response-related activity supports the idea that the anterior MTL is involved in the conceptual category representations of visual scenes^8,15,27–29^. Notably, anterior MTL HFA tracked scene category responses in both subsequent hit and miss trials, indicating that while it reflects category processing, it may not be the primary feature supporting memory formation in the anterior MTL. These latency findings support the hierarchical processing along visual pathways^88–90^, in which perceptual details are represented earlier in more posterior regions and stimulus identity representations are formed over time in anterior regions.

### Posterior MTL HFA and memory

Extending the widely observed preferential memory responses to visual scenes in the posterior MTL in fMRI studies^9,22,23,33,34^, we observed increased HFA in subsequent hit trials (i.e., positive subsequent memory effects) after scene onset and before subjects’ scene category responses. In addition, increased HFA before subjects’ scene category responses was positively associated with memory performance, underscoring HFA as a neurophysiological feature of the posterior MTL supporting memory. In the anterior MTL, we observed positive HFA subsequent memory effects after scene onset and before scene category responses. However, unlike the pronounced HFA increases in the posterior MTL, the anterior MTL exhibited an overall reduced level of HFA and neither of the subsequent memory effects clusters was associated with memory performance. The reduced level of HFA in the anterior MTL might be due to a smaller number of neurons coding conceptual information^90,91^. Future studies incorporating human single-neuron recordings are needed to investigate the neural computation underlying conceptual representation in the anterior MTL^90,92^.

### Anterior MTL theta oscillation and memory

By systemically detecting and parameterizing neural oscillations in the posterior and anterior MTL, we confirmed that theta oscillations, ranging from ∼2-10 Hz, dominate the developing MTL and exhibit significant inter-individual variability. This observation supports the use of individually identified theta oscillations, rather than the canonical broadband theta range (∼4-8 Hz), to accurately assess individual differences in neural oscillations and their functional roles^36,80^. After parameterizing theta oscillations by center frequency, power, and bandwidth, we examined the differences in these parameters between subsequent hit and miss trials and the posterior and anterior MTL. Except for theta power differences between subsequent hit and miss trials, theta parameters did not differ in relation to subsequent memory outcome.

We observed theta power decreases in subsequent hit trials in both the posterior and anterior MTL, consistent with known broadband low-frequency power decreases during successful memory formation^36,40^ and extending the idea of cortical desynchronization in the service of information representation for successful memory^93^ to individually detected narrowband low-frequency oscillations. The lack of subsequent memory effects in theta frequency and bandwidth implies that theta oscillations provide an infrastructure for memory processes^36^, with the difference between successful and unsuccessful memory formation lying in theta-associated mechanisms, rather than in the oscillations themselves.

Although no regional differences in theta parameters were observed, the posterior and anterior MTL differed in how theta parameters were associated with memory performance and age. In the anterior MTL, we observed that faster theta frequency was associated with better memory performance. In line with the correlation between theta frequency and spatial navigation speed in the hippocampus^94,95^, increased frequency may underlie increased information processing^40,96^, potentially due to faster fluctuations between neuronal excitation and inhabitation^58^. Future studies that detect and parameterize theta during different cognitive tasks, as well as employ computational models, are needed to further elucidate the underlying mechanisms linking theta frequency to cognitive performance. In the posterior MTL, we observed an age-related increase in theta power. Because cortical theta power decreases are associated with memory, the age-related increase in theta power may not be related to memory and rather reflect aspects of posterior MTL development^33,35^ that can be elucidated in further research.

### Theta-HFA coupling in the anterior MTL supports memory

Theta oscillations are posited to underlie memory by facilitating information integration^55–58^. Based on the temporal relationship between neuronal spiking and ongoing theta oscillations in the rodent hippocampus^97^, theta is posited to organize neuronal spiking and, by extension, gamma activity or HFA representing information within each cycle^55–58^. In line with the hypothesized functional role of theta, we observed increased theta-HFA coupling in the anterior MTL during successful memory formation, extending the theta-gamma code in the hippocampus^98,99^ and neocortex^100^ to theta-HFA coupling in the MTL cortex. This memory-related theta-HFA coupling occurred before subjects’ scene category responses, indicating that conceptual category representation in the anterior MTL may be supported through the integration of visual information facilitated by theta-HFA coupling. The theta-HFA coupling subsequent memory effects were not associated with individual differences in memory performance. This is in contrast to theta frequency in the anterior MTL, which was related to individual differences in memory performance, raising the possibility that faster theta serves as a more finely tuned mechanism supporting memory^36,42^ and that its specific frequency is more relevant to memory outcome than theta-HFA coupling strength.

### Posterior-anterior theta coupling supports memory

Theta oscillations underlie information transfer between brain regions via phase coupling^59,61,62^. Consistent with previous findings of increased inter-regional theta coupling during successful memory formation^36,101,102^, we observed increased posterior-anterior MTL theta phase coupling during successful memory formation, likely indicating information transfer between the posterior and anterior MTL. We further explored the interaction between the posterior and anterior MTL by examining the inter-regional cross-frequency phase-amplitude coupling, an inter-regional interaction mechanism that has been reported within hippocampal subfields and between hippocampus and extra-hippocampal cortex in support of memory formation^82,84,103,104^. Specifically, we examined whether posterior MTL HFA, which we identified as relevant to memory, is coupled to anterior MTL theta, which we also identified as relevant to memory. We did not observe anterior-posterior theta-HFA coupling subsequent memory effects in either stimulus-locked or response-locked data (see Fig. S3), suggesting that distinct, rather than directly coupled, neurophysiological features support memory along the MTL long axis.

However, because of the theta coupling between the posterior and anterior MTL, we further explored the relationship between posterior MTL HFA and anterior MTL theta by directly correlating these measures. We observed that increased posterior HFA was associated with increased anterior theta frequency. Moreover, this positive correlation was specific to subsequent hit but not miss trials for response-related posterior HFA subsequent memory effects (Fig. 7c, right). Posterior MTL HFA was not correlated with other anterior theta parameters (i.e., power, bandwidth; see Fig. S4), suggesting a selective relationship between posterior MTL HFA and anterior MTL theta frequency.

These findings indicate increased information transfer and processing along the MTL long axis in support of memory, and that posterior-anterior theta coupling may link distinct neurophysiological features across regions to support the multifaceted representations of visual scenes.

### Age-invariant neurophysiological features in the posterior and anterior MTL

Similar to the content-specific organization of the MTL long axis in adults^7–9,22–27^, the observed distinct activity latencies and neurophysiological features suggest different representations along the long axis of the developing MTL. Moreover, consistent with age-invariant memory-related activation reported in fMRI studies^31,32^, memory-related effects across the examined neurophysiological features in either the posterior or anterior MTL did not differ by age. The similarity of representation patterns to those in adults and the absence of age differences in memory-related neurophysiological features suggest early development of the MTL^3,4,31,105^.

However, some age differences point to ongoing development, particularly of the posterior MTL. The age-related increase in posterior MTL theta power, considered together with subsequent memory theta power decreases, suggests a reduced reliance on theta oscillations across development. We speculate that this reduced involvement of theta is likely due to the increased specialization of scene perception in the posterior MTL with development^33–35,106^.

Age invariance in memory-related effects may also reflect task or stimulus characteristics. Subjects were asked to study complex visual scenes presented on the screen for 3 seconds and their memory was probed with a recognition test. This task was selected because it involves simple instructions and minimized demands that greatly differ by age such as reading fluency. Future developmental intracranial EEG studies that employ diverse memory tasks can corroborate the findings across different stimulus categories^34^, such as scenes, objects, faces, and words and with various mnemonic difficulty levels^35,106^.

An important additional consideration when investigating age-related differences in regional brain effects is that development likely manifests as the maturation of connectivity between these regions and other key memory brain regions. Previous studies have shown differential connectivity patterns along the MTL long axis. For example, the posterior MTL primarily connects with the posterior hippocampus and other regions within the default mode network, such as the medial parietal cortex and precuneus, while the anterior MTL primarily connects with the anterior hippocampus, amygdala, and orbitofrontal cortex^8,10–12,107^. Differential connectivity may serve as the infrastructure for functional specialization along the MTL long axis. Future studies investigating the development of connectivity mechanisms are needed to better understand the development of the functional specialization of the MTL.

### Limitations and future directions

Developmental intracranial EEG offers an unparalleled opportunity to investigate the fundamental mechanisms of brain development underlying cognitive development, as it provides localized and temporally precise measures of brain activity. However, such data are rare, and the small sample sizes typically available for evaluating individual differences^36,37,40^ may limit our ability to detect potential developmental variability in neurophysiological features^41,42,63^. This limitation necessitates cautious interpretation of the age-invariant findings. Increasing sample size through multi-site collaboration and data sharing is recommended to overcome the sample size issues in developmental intracranial EEG studies^41,42,63^. We also note that, although the task used to probe memory is suitable for both children and adults and is commonly used in developmental research, it uses only visual scenes and incorporates relatively simple memory demands. Employing a task that directly manipulates content-specific representations, or distinguishes between perceptual and semantic processing, would likely help identify more nuanced effects or differential content representation along the long axis of the developing MTL.

We also note that we employed a pragmatic, convenience-based distinction to classify electrodes as belonging either to the anterior or posterior MTL; this approach was largely agnostic to known anatomically defined regions, namely the parahippocampal, entorhinal, and perirhinal cortices. This choice was driven by practical limitations, including the absence of harmonized protocols for segmenting MTL subregions, substantial individual variability in anatomical landmarks, and the requirement for high-resolution imaging data^108,109^. We confirmed, however, that the midpoint between the most anterior and most posterior electrodes (y = -14.35 mm) used to classify posterior and anterior MTL electrodes is consistent with previous fMRI studies that separate parahippocampal and perirhinal cortices^11,12^. Nonetheless, we referred to the posterior and anterior MTL, rather than to the parahippocampal and perirhinal cortices, to be consistent with our classification approach. Future studies with high-resolution structural MRI data and manual demarcation of MTL subregions at the individual level^22,26,110–112^ can further probe anatomical specificity in the neurophysiological features. Lastly, the data presented in the current study are from surface electrodes located on the MTL cortex. The hippocampus exhibits a representation gradient along its long axis^113^, and future research incorporating hippocampal recordings from stereoencephalography (sEEG) depth electrodes would enable a more comprehensive examination of MTL structures. Such investigation, combining MTL surface electrodes with hippocampus depth electrodes, could provide valuable insights into how the hippocampus modulates content representations in the neocortex^8^.

### Conclusions

The developing MTL is differentially involved in representations of perceptual and conceptual information of visual scenes along its posterior-to-anterior axis, as evidenced by distinct activity latencies and neurophysiological features. With the unparalleled spatiotemporal resolution afforded by ECoG, we show that the timing of HFA tracks scene onset in the posterior MTL and subjects’ scene category responses in the anterior MTL. Our findings further show that, in the posterior MTL, HFA predicts successful memory formation and is positively linked to memory performance. In contrast, in the anterior MTL, theta frequency is linked to memory performance, and theta-HFA phase-amplitude coupling before subjects’ scene category responses predicts successful memory formation. Collectively, our findings reveal distinct neurophysiological mechanisms along the MTL long axis that support the multifaceted representations of episodic memory formation in children and adolescents.

## MATERIALS AND METHODS

### Method details

#### Pediatric patients

In total, 23 neurosurgical patients (12 females and 11 males; 5.9-20.5 years of age; M ± SD, 13.4 ± 4.0 years; Table S1) from the Children’s Hospital of Michigan with subdural electrodes implanted for the clinical evaluation of refractory epilepsy participated in this study. Written informed consent was obtained from patients aged 18 years and older and from the guardians of subjects under 18 years. Written assent was obtained from subjects aged 13-17 years, and oral assent was obtained from younger children. This study was approved by the Institutional Review Board at Wayne State University in accordance with the Declaration of Helsinki. Among these patients, 17 patients (8 females and 9 males; 6.2-20.5 years of age; M ± SD, 13.3 ± 3.7 years) with electrode coverage in the posterior MTL, 20 patients (10 females and 10 males; 5.9-19.3 years of age; M ± SD, 13.0 ± 3.9 years) with electrode coverage in the anterior MTL, and 14 patients (6 females and 8 males; 6.2-17.1 years of age; M ± SD, 12.6 ± 3.4 years) with electrode coverage in both the posterior and anterior MTL.

#### Behavioral task

Subjects completed a scene recognition memory task (Fig. 1a) that has been used to study the neurophysiological mechanisms of memory with pediatric samples^36,37,40,114,115^. Pictures of scenes were presented in consecutive study-test cycles. In each study cycle, subjects studied 40 scenes of equal number of indoor and outdoor scenes, each presented 3 s following a 0.5-s fixation screen. Subjects verbalized whether each scene was “indoor” or “outdoor”. Responses were coded as correct or incorrect via offline review of individual audio recordings. Analysis of electrophysiological data was restricted to studied trials with correct indoor/outdoor responses. In each test cycle, subjects were tested on all 40 studied scenes, intermixed with 20 new scenes as foils. Subjects were instructed to respond “old” if they remembered seeing the scene and “new” if they thought they had not seen the scene in the study cycle. Each scene remained on the screen until a new/old response was given, following a 0.5-s fixation screen. Responses were coded as hit (“old” response to studied scene), miss (“new” response to studied scene), correct rejection (“new” response to foil), or false alarm (“old” response to foil). Memory performance was measured with recognition accuracy, which was calculated by hit rate (number of hits out of the total number of studied scenes) minus false alarm rate (number of false alarms out of total number of foils). Electrophysiological data for studied scenes were analyzed as a function of subsequent memory at test (i.e., subsequent hit or miss)^36,37,40^. Across all 23 subjects, 43% (n = 10) completed one study-test cycle, and 57% (n = 13) completed two study-test cycles.

#### Electrode placement and localization

Platinum macro-electrodes (4 mm diameter, 10 mm intercontact distance) were surgically implanted for extra-operative ECoG monitoring based solely on the clinical needs of each patient. Three-dimensional electrode reconstructions were created by co-registering post-implantation planar x-ray images of the cortical surface with preoperative T1-weighted spoiled gradient-echo magnetic resonance images^64^.

Automatic parcellation of cortical gyri was performed using FreeSurfer software^116^ and anatomical labels were assigned to each electrode site^117^. Subjects were included in this study by artifact-free electrode coverage of the MTL, as determined by review of individual reconstructions and automatic parcellation results. Electrodes were plotted on MNI space for visual representation across all subjects (Fig. 2c) using the Fieldtrip toolbox^118^ for MATLAB (MathWorks Inc., Natick, MA).

#### ECoG data acquisition and preprocessing

ECoG data were acquired using a 192-channel Nihon Koden Neurofax 1100A Digital System, sampled at 1kHz. Raw electrophysiological data were filtered with 0.1-Hz high-pass and 300-Hz low-pass finite impulse response filters, and 60-Hz line noise harmonics were removed using discrete Fourier transform. Continuous study data were demeaned, epoched into 4.5-s trials (-1 to 3.5 s from scene onset), and manually inspected blind to electrode locations and experimental task parameters. Electrodes overlaying seizure onset zones^119^ and electrodes and epochs displaying epileptiform activity or artifactual signal (from poor contact, machine noise, etc.) were excluded.

Each artifact-free electrode was re-referenced to the common average of all artifact-free electrodes. The re-referenced data were re-inspected to reject any trials with residual noise. The final data set included a median(IQR) of 4(2) (range, 1 to 10) artifact-free and nonpathological MTL electrodes and a mean ± SD of 48 ± 17 (22 to 76) study trials per subject.

### Neurophysiological data analysis

#### High-frequency activity analysis

Time-frequency representations of high-frequency activity (HFA, 70-150 Hz) were quantified using a multitapering approach^120^ with the FieldTrip toolbox^118^. Study trials were zero-padded to the next power of 2 to minimize edge artifacts. The multitaper frequency spectrum was then calculated by sliding a 0.25-s window in 0.025-s steps across 30 logarithmically spaced, partially overlapping frequency bands centered from 70 to 150 Hz, with ½ fractional bandwidth at each center frequency.

The frequency spectrum of encoding interval (-0.15 to 3 s from scene onset) was corrected on the pre-stimulus baseline interval (-0.45 to -0.15 s from scene onset) with statistical bootstrapping. For each electrode and frequency, data from baseline interval were pooled across all trials from which r data points (r = number of trials in that subject’s data) were randomly selected and averaged. This process was repeated 1000 times to create normal distributions of the baseline data. Power from encoding interval was z-scored per time point on the pre-stimulus baseline distributions. This procedure adjusts the time-frequency representations to correct for 1/f power scaling and reveals event-related spectral activity^36,37,40,68,121^. The mean of the z-scored power across frequencies was then taken as a single HFA time series for future analysis. The output of HFA analysis was initially time-locked to scene onset. The response-locked HFA was obtained by aligning the z-scored power with each trial’s RT, setting the RT as time zero. Subsequent memory analysis on HFA was conducted on both the stimulus-locked (-0.15 to 1 s) and response-locked (-1 to 0.2 s) data.

HFA peak latency was analyzed on the single-trial level. Each HFA time series was smoothed using a moving average with a window size of 0.05 s. HFA peak latency was defined as the time point of the maximum HFA between scene onset and 0.2 s after scene category response onset. The peak latency of each electrode was calculated by averaging latencies across subsequent hit or miss trials for statistical modeling.

#### Oscillation detection

SpecParam was used to isolate the oscillatory component from the aperiodic 1/f component^80^ between 1-30 Hz. For each electrode, the power spectrum of study data was calculated with Fieldtrip toolbox^118^ and submitted to SpecParam for oscillation detection. The power spectrum was calculated with fast Fourier transform for each study trial from -0.45 to 3 s from scene onset with 2-s segment and 1.5-s overlap. Each segment was zero-padded to the next power of 2 and tapered with a Hanning window to minimize edge effects. The mean of the segments power spectra was then taken as each trial’s power spectrum. The power spectra of hit and miss trials were then averaged separately for oscillation detection.

SpecParam models the power spectrum by initially fitting a log-logistic function to capture the aperiodic component. Once parameterized, the aperiodic component is subtracted from the power spectrum. The remaining data is then fitted with Gaussian functions iteratively to identify and parameterize oscillatory peaks (Fig. 5a) until either no peaks meet the predefined threshold or the desired number of peaks is reached. Each peak is parameterized with center frequency, power, and bandwidth. The SpecParam algorithm used with settings of {peak_width_limits = [2,6], max_n_peaks = 6, min_peak_height = 0.1, peak_threshold = 1.5, aperiodic_mode = ‘fixed’}. The peak with the largest power between 1-30 Hz was chosen to examine the center frequency distribution. After confirming the largest oscillations clustered within the theta range (Fig. 5c), the largest peak between 1-10 Hz was chosen as theta oscillation for each electrode. The parameterization procedure was done separately for subsequent hit and miss trials.

#### Phase-amplitude coupling

Intra-regional and inter-regional event-related phase-amplitude coupling (ERPAC) was analyzed to capture the transient and temporal dynamics of phase-amplitude coupling^122^. Study trials were padded with reflection padding and then bandpass filtered at the detected theta center frequency and bandwidth and high-frequency band (70-150 Hz) for each electrode, separately for subsequent hit and miss trials, following the removal of ERP^36,121,123^. The Hilbert transform was then used to extract phase values from the theta time series and amplitude values from the high-frequency time series. To evaluate the dependence of high-frequency amplitude on theta phases, the high-frequency amplitude was bandpass filtered a second time at the detected theta frequency and bandwidth. Phase values were extracted from this filtered signal using the Hilbert transform. Phase-locking value (PLV) was then calculated as a measure of ERPAC to quantify the dependence of high-frequency amplitude on theta phases^36^. For inter-regional ERPAC analysis, the anterior MTL is the phase providing region and the posterior MTL is the amplitude providing region. Inter-regional ERPAC was analyzed between all possible anterior-posterior pairs for each subject.

PLV was calculated as the consistency of phase differences at each time point between theta phases and the phases of theta-filtered high-frequency amplitude separately across subsequent hit and miss trials. To account for differences in trial counts between subsequent hit and miss trials, the raw PLV was normalized using statistical bootstrapping. A surrogate PLV distribution was created by randomly shuffling trial labels of theta phases and recalculating the PLV 1,000 times. The raw PLV was then z-scored against the surrogate distribution^36^. ERPAC analysis was conducted on data time-locked to scene onset (-1 to 3.5 s from scene onset) and to scene category response onset (-1.25 to 0.25 s from response onset). Subsequent memory analysis was conducted on the normalized PLV within the time window of interest (stimulus-locked: -0.2 to 1 s from scene onset; response-locked: -1 to 0.2 s).

To identify the theta phase associated with increased high-frequency amplitude in the anterior MTL, the preferred phase analysis was conducted during the time window of significant SMEs in ERPAC, extending by ± 0.2 s (see Fig. 6d). For each trial, theta phases were divided into 18 bins ranging from −π to π. High-frequency amplitude was binned based on the theta phases and normalized to the percentage of the largest binned amplitude.

#### Inter-regional phase coupling

Study trials were padded with reflection padding and then bandpass filtered at the detected theta center frequency and bandwidth of each electrode, following the removal of ERP^36,121,123^. The Hilbert transform was then used to extract phase values from the theta time series. PLV was calculated between all posterior-anterior MTL electrode pairs as a measure of phase coupling. Here, PLV quantifies the consistency of phase differences between each electrode pair at each time point separately across subsequent hit and miss trials. To account for differences in trial counts between subsequent hit and miss trials, the raw PLV was normalized using statistical bootstrapping. For each electrode pair, a surrogate PLV distribution was created by randomly shuffling trial labels in one electrode and recalculating the PLV 1,000 times.

The raw PLV was then z-scored against the surrogate distribution^36^. Phase coupling analysis was conducted on data time-locked to scene onset (-1 to 3.5 s from scene onset) and to scene category response onset (-1.25 to 0.25 s from response onset). Subsequent memory analysis was conducted on the normalized PLV within the time window of interest (stimulus-locked: -0.2 to 1 s from scene onset; response-locked: -1 to 0.2 s).

### Statistical analysis

#### HFA peak latency

To model HFA peak latency differences along the long axis of MTL, a linear mixed-effects (LME) model with subsequent memory (i.e., hit vs. miss) and electrodes’ y-coordinates as fixed effects and subjects and nested electrodes as random effects was used. A significant subsequent memory × y-coordinates interaction effect was followed up with separate linear mixed-effects models for hit and miss, with y-coordinates as fixed effects and subjects and nested electrodes as random effects.

To model the correlation of HFA peak latency with subjects’ indoor/outdoor RT and to examine if the correlation differs by region of interest (ROI; posterior vs. anterior MTL) and subsequent memory (hit vs. miss), a linear mixed-effects model with ROI, subsequent memory, and RT as fixed effects and subjects and nested electrodes as random effects was used. A significant subsequent ROI × subsequent memory × RT interaction effect was followed up with separate linear mixed-effects models for posterior and anterior MTL, with subsequent memory and RT as fixed effects and subjects and nested electrodes as random effects. To follow up on the significant subsequent memory × RT interaction in the posterior MTL, additional models for subsequent hit and miss were conducted with RT as fixed effect and subjects and nested electrodes as random effects.

#### Theta parameters

After detecting and parameterizing theta oscillations between 1-10 Hz for each electrode, LME models were used to test if theta parameters (i.e., theta frequency, power, and bandwidth) differ by regions and subsequent memory. For each theta parameter, a LME model was conducted with regions and subsequent memory as fixed effects and subjects and nested electrodes as random effects.

To examine the individual differences in theta parameters for each region, LME models were conducted separately for the posterior and anterior MTL. For each theta parameter, LME was run with subsequent memory, recognition accuracy, and age as fixed effects and subjects and nested electrodes as random effects.

#### Subsequent memory effects (SMEs)

SMEs were analyzed on HFA, phase-amplitude coupling, and phase coupling data to test the differences between subsequent hit and miss trials. For each region and measure, subsequent memory analysis was conducted with an LME model and cluster-based correction was used to correct for multiple comparisons. LME was conducted for each time point with subsequent memory as fixed effects and subjects and nested electrodes as random effects. To correct for multiple comparisons, the hit and miss trial labels were randomly shuffled within each electrode and the model was re-run. This procedure was repeated 1,000 times. The size of the maximum cluster of significant SMEs (p < 0.05) from each run was extracted to create the random permutation distribution. The cluster size of the observed SMEs (p < 0.05) was compared against the random permutation distribution to obtain the corrected p-value of each subsequent memory cluster^36,124^. SMEs that survived the cluster-based correction (corrected p < 0.05) were reported. To disentangle the scene-induced and response-related effects, the subsequent memory analysis was conducted on the data locked to scene onset and locked to scene category response onset.

#### Inter-individual differences in subsequent memory effects

Multiple regression analyses were conducted with recognition accuracy and age as predictors to investigate behavioral relevance and age-related differences of the SMEs^36,40^. The differences between subsequent hit and miss were averaged across electrodes for each subject and across the time window of each significant subsequent memory cluster (corrected p < 0.05) and submitted to multiple regression with recognition accuracy and age as predictors.

#### Correlation between posterior MTL HFA and anterior MTL theta parameters

The Pearson correlation between posterior MTL HFA and anterior MTL theta parameters (i.e., frequency, power, and bandwidth) was calculated to assess the association between the distinct neurophysiological features in the posterior and anterior MTL. Posterior HFA was obtained separately for subsequent hit and miss trials by averaging HFA within the time window of significant SMEs (Fig. 4a). The Pearson correlation was then calculated between the posterior HFA and the anterior MTL theta parameters separately for subsequent hit and miss trials.

## Data Availability

De-identified raw data are deposited to the NIMH public database (DOI pending acceptance). Additional data related to the paper may be requested from the authors.

## Code Availability

Custom code used for the analyses in this study is available at GitHub: https://github.com/qinyin07/mtl_long_axis.

## Acknowledgements

We thank the subjects and families who participated in this study. We thank J. S. Damoiseaux, A. M. Daugherty, T. Fischer, R. Homayouni, K. L. Canada, and Z. Chen for support. We thank Y. Nakai, M. Sonoda, N. Kuroda, A. T. Shafer, M. M. Malik, and Y. Chen for assistance. This work was supported by funding from the National Institutes of Health (R01MH107512 to NO and R00NS115918 to ELJ).

## Author Contributions

Conceptualization: Q.Y., N.O., and E.L.J.; Methodology: N.O., Q.Y., and E.L.J.; Software: Q.Y., E.L.J., and A.J.D.; Validation: Q.Y., N.O., and E.L.J.; Formal analysis: Q.Y.; Investigation: N.O., E.A., E.L.J., and R.T.K.; Resources: N.O. and E.L.J.; Data curation: Q.Y., E.L.J., and N.O.; Visualization: Q.Y. and E.L.J; Writing - original draft: Q.Y.; Writing - review and editing: Q.Y., N.O., E.L.J., R.T.K., and A.J.D.; Supervision: N.O.; Project administration: N.O., E.L.J., and Q.Y.; Funding acquisition: N.O., E.L.J., R.T.K., and E.A.

## Competing Interests

None.

## Notes

### Competing Interest Statement

The authors have declared no competing interest.

