## supplemental figures and tables for "Distinct neurophysiological features and memory representations along the long axis of the developing medial temporal lobe"

### Supplementary materials

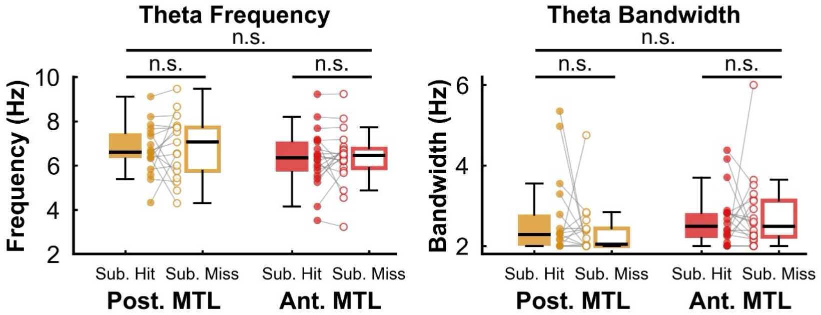


**Figure S1. Theta frequency and bandwidth between the posterior and anterior MTL, and between subsequent hit and miss trials.** Sub. Hit, subsequent hit; Sub. Miss, subsequent miss; Post. MTL, posterior MTL; Ant. MTL, anterior MTL.

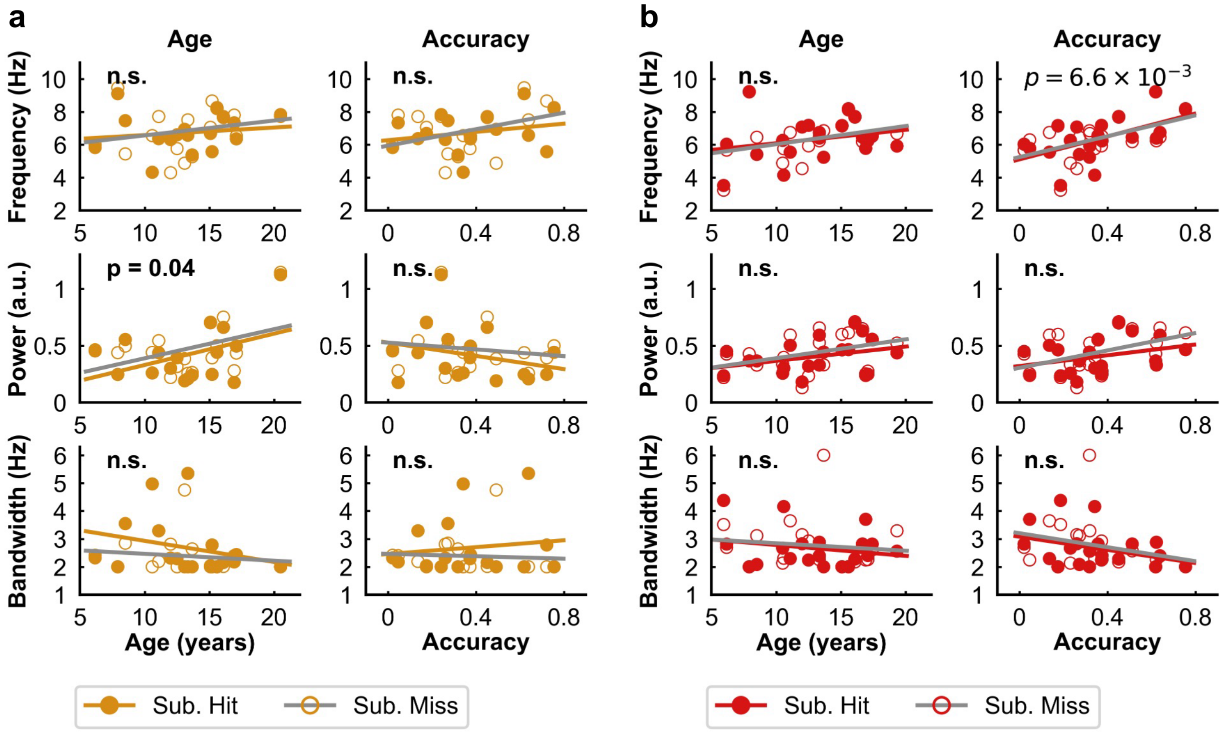


**Figure S2. Individual differences in theta parameters.**
(**a**) The association of theta parameters with age and memory performance in the posterior MTL. From left to right: age effect and recognition accuracy effect. From top to bottom: theta frequency, theta power, and theta bandwidth. n.s., p > 0.05. Sub. Hit, subsequent hit; Sub. Miss, subsequent miss.
(**b**) Same as (a) for theta parameters in the anterior MTL.

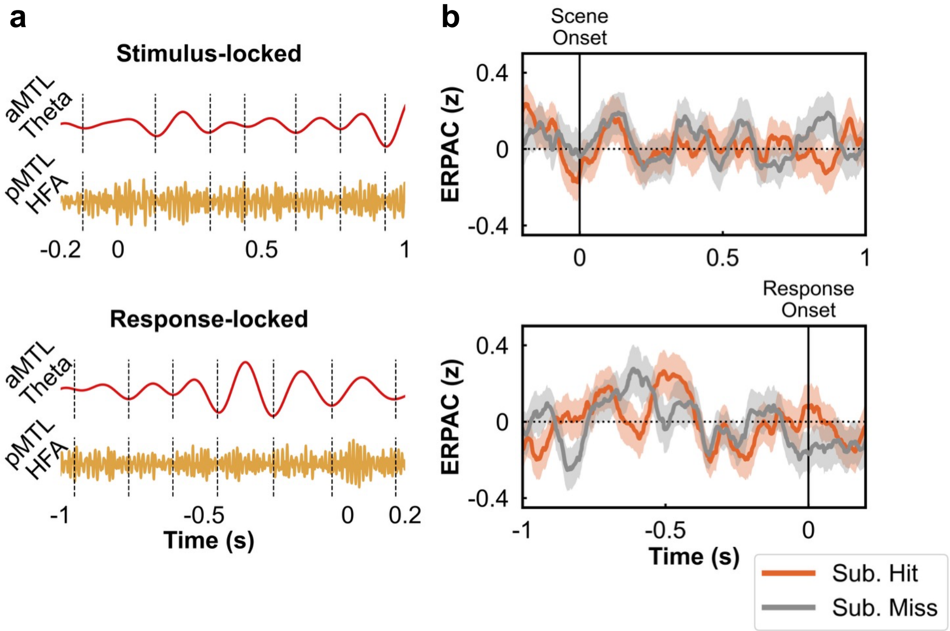


**Figure S3. Posterior-anterior inter-regional ERPAC.**
(**a**) Top: theta-filtered signal of a representative subsequent hit trial from an anterior MTL electrode (top), and HFA-filtered signal of the same trial from a posterior MTL electrode for stimulus-locked data. Bottom: The same signal locked to scene category response onset. Dashed vertical lines indicate theta troughs.
(**b**) Anterior-posterior event-related phase-amplitude coupling (ERPAC) in stimulus-locked (top) and response-locked (bottom) data. Shaded areas represent the standard error of the mean across subjects. No significant SMEs were observed for inter-regional ERPAC. Sub. Hit, subsequent hit; Sub. Miss, subsequent miss.

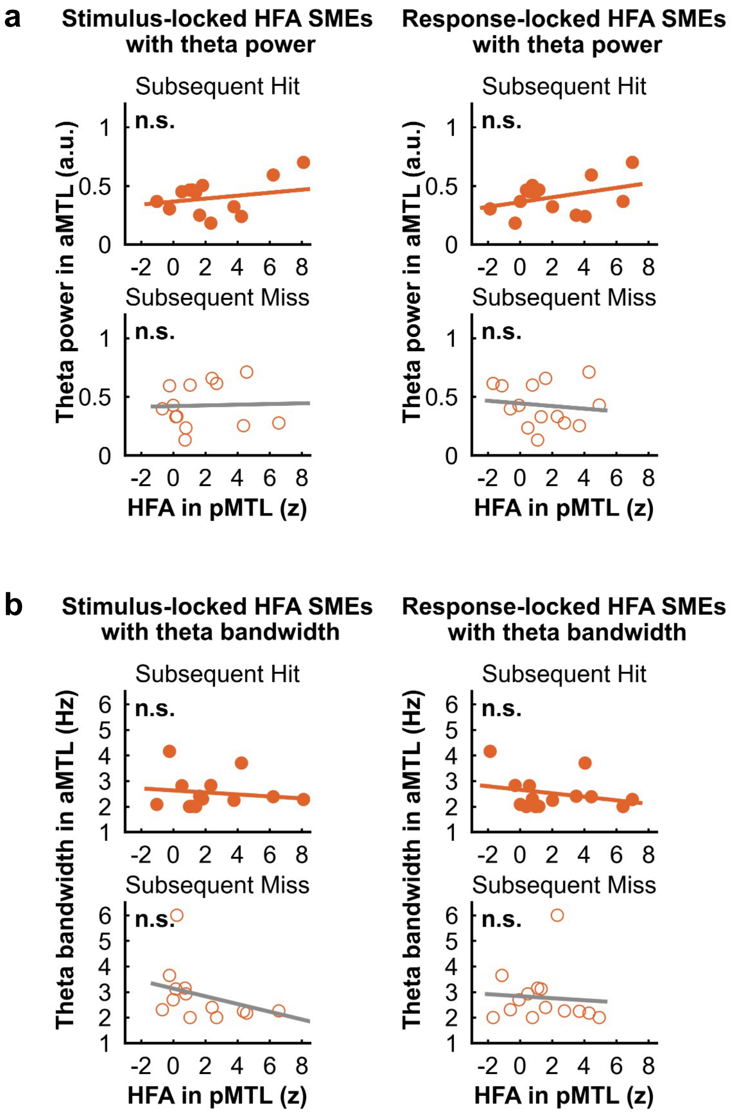


**Figure S4. Correlation of posterior MTL HFA with anterior theta power and bandwidth.** (**a**) Correlation of posterior MTL HFA with anterior theta power. Left: HFA from the time window of significant SMEs from stimulus-locked data, see Fig. 4a, top, shaded area. Right: HFA from the time window of significant SMEs from response-locked data, see Fig. 4a, bottom, shaded area (**b**) Same as (a) for theta bandwidth in the anterior MTL. n.s., $p > 0.05$.

**Table S1. Subject information.**

| Subject | Age (years) | Sex | Study trials | Study accuracy | Test accuracy | Seizure Onset | Post. MTL | Ant. MTL |
|---|---|---|---|---|---|---|---|---|
| S1 | 13.67 | F | 81 | 0.94 | 0.32 | Lt T | 1 | 1 |
| S2 | 12.00 | F | 80 | 0.89 | 0.26 | Lt FP | 2 | 2 |
| S3 | 15.08 | M | 40 | 1.00 | 0.18 | Rt F | 2 | 2 |
| S4 | 12.50 | F | 80 | 0.93 | 0.37 | Rt P | 2 | 3 |
| S5 | 10.50 | F | 80 | 0.96 | 0.23 | Rt T | 0 | 4 |
| S6 | 15.58 | M | 70 | 0.99 | 0.76 | Lt T | 1 | 2 |
| S7 | 10.58 | M | 40 | 0.95 | 0.34 | Rt P | 1 | 6 |
| S8 | 16.92 | M | 80 | 0.96 | 0.05 | NA | 2 | 2 |
| S9 | 6.17 | F | 80 | 0.95 | 0.02 | Lt PO | 3 | 7 |
| S10 | 11.08 | M | 80 | 0.94 | 0.14 | Rt F & Lt F | 3 | 2 |
| S11 | 19.33 | F | 80 | 0.95 | 0.32 | Lt F | 0 | 3 |
| S12 | 16.08 | F | 80 | 0.95 | 0.45 | Rt P | 4 | 4 |
| S13 | 8.50 | M | 40 | 0.83 | 0.27 | Lt P | 1 | 2 |
| S14 | 5.94 | F | 40 | 0.98 | 0.19 | Rt F | 0 | 3 |
| S15 | 16.67 | F | 80 | 1.00 | 0.51 | NA | 0 | 3 |
| S16 | 13.33 | M | 40 | 0.90 | 0.64 | Rt F | 1 | 4 |
| S17 | 13.08 | F | 40 | 0.95 | 0.49 | Rt TPO | 1 | 0 |
| S18 | 20.50 | M | 80 | 0.95 | 0.24 | Rt TF | 1 | 0 |
| S19 | 15.21 | F | 40 | 0.98 | 0.72 | Rt TPO | 1 | 0 |
| S20 | 17.42 | M | 40 | 0.88 | 0.36 | Lt F | 0 | 2 |
| S21 | 13.33 | M | 80 | 0.99 | 0.62 | Rt F | 0 | 3 |
| S22 | 7.92 | F | 40 | 0.98 | 0.62 | NA | 2 | 2 |
| S23 | 17.08 | M | 40 | 1.00 | 0.38 | NA | 1 | 4 |

*Study accuracy* indicates indoor/outdoor response accuracy. *Test accuracy* indicates recognition accuracy, i.e., hit rate – false alarm rate. *Post. MTL* and *Ant. MTL* indicates the number of electrodes per region of interest. Lt = left; Rt = right; F = frontal lobe; T = temporal lobe; P = parietal lobe; O = occipital lobe; NA = seizures were not captured; Post. = posterior; Ant. = anterior.

**Table S2. LME model estimate for region, subsequent memory, and RT effects on HFA peak latency.**

| Variables | Beta | Std. Error | t | P-value |
|---|---|---|---|---|
| Region (post. vs. ant.) | -0.38 | 0.20 | -1.89 | 0.06 |
| Subsequent memory (hit vs. miss) | 0.11 | 0.13 | 0.84 | 0.40 |
| RT | 0.27 | 0.21 | 1.29 | 0.20 |
| Region × Subsequent memory | 0.17 | 0.13 | 1.31 | 0.19 |
| Region × RT | -0.41 | 0.19 | -2.08 | 0.04* |
| Subsequent memory × RT | 0.17 | 0.13 | 1.37 | 0.17 |
| Region × Subsequent memory × RT | 0.27 | 0.12 | 2.22 | 0.03* |

* p< 0.05; post., posterior MTL; ant., anterior MTL

**Table S3. Post hoc per region LME models estimate for subsequent memory and RT effects on HFA peak latency.**

| Region | Variables | Beta | Std. Error | t | P-value |
|---|---|---|---|---|---|
| Posterior MTL | Subsequent memory (hit vs. miss) | 0.27 | 0.15 | 1.76 | 0.08 |
| | RT | -0.22 | 0.25 | -0.88 | 0.38 |
| | Subsequent memory × RT | 0.48 | 0.14 | 3.32 | $1.6 \times 10^{-3}$ ** |
| Anterior MTL | Subsequent memory (hit vs. miss) | -0.06 | 0.14 | -0.44 | 0.66 |
| | RT | 0.74 | 0.24 | 3.04 | $2.9 \times 10^{-3}$ ** |
| | Subsequent memory × RT | -0.12 | 0.16 | -0.75 | 0.46 |

**p < 0.01

**Table S4. Multiple regression model estimate for age and recognition accuracy effects on HFA subsequent memory effects.**

| Region | Referenced time | Variables | Beta | Std. Error | t | P-value |
|---|---|---|---|---|---|---|
| Posterior MTL | Scene-induced SME | Age | -0.16 | 0.3 | -0.55 | 0.59 |
| | | Accuracy | -0.05 | 0.27 | -0.18 | 0.86 |
| | | Age × Accuracy | -0.19 | 0.28 | -0.67 | 0.51 |
| | Response-related SME | Age | 0.13 | 0.24 | 0.53 | 0.60 |
| | | Accuracy | 0.57 | 0.22 | 2.57 | 0.02* |
| | | Age × Accuracy | 0.22 | 0.23 | 0.97 | 0.35 |
| Anterior MTL | Scene-induced SME | Age | 0.06 | 0.27 | 0.24 | 0.81 |
| | | Accuracy | 0.16 | 0.25 | 0.65 | 0.52 |
| | | Age × Accuracy | -0.09 | 0.27 | -0.33 | 0.75 |
| | Response-related SME | Age | 0.34 | 0.26 | 1.33 | 0.20 |
| | | Accuracy | -0.13 | 0.24 | -0.52 | 0.61 |
| | | Age × Accuracy | 0.20 | 0.26 | 0.78 | 0.45 |

* $p < 0.05$

**Table S5. LME model estimate for region and subsequent memory differences on oscillation parameters.**

| Parameters | Variables | Beta | Std. Error | t | P-value |
|---|---|---|---|---|---|
| Theta Frequency | Region (post. vs. ant.) | 0.08 | 0.06 | 1.33 | 0.18 |
| | Subsequent memory (hit vs. miss) | -0.02 | 0.05 | -0.45 | 0.65 |
| | Region × Subsequent memory | -0.02 | 0.05 | -0.37 | 0.72 |
| Theta Power | Region (post. vs. ant.) | 0.04 | 0.05 | 0.96 | 0.34 |
| | Subsequent memory (hit vs. miss) | -0.11 | 0.04 | -2.95 | $3.7\times10^{-3}$** |
| | Region × Subsequent memory | -0.02 | 0.04 | -0.42 | 0.68 |
| Theta Bandwidth | Region (post. vs. ant.) | -0.11 | 0.08 | -1.44 | 0.15 |
| | Subsequent memory (hit vs. miss) | 0.12 | 0.07 | 1.57 | 0.12 |
| | Region × Subsequent memory | 0.02 | 0.07 | 0.28 | 0.78 |

** $p < 0.01$

**Table S6. LME model estimate for individual differences on posterior MTL theta parameters.**

| Parameters | Variables | Beta | Std. Error | t | P-value |
|---|---|---|---|---|---|
| Theta Frequency | Age | 0.05 | 0.18 | 0.29 | 0.77 |
| | Accuracy | 0.30 | 0.16 | 1.81 | 0.08 |
| | Subsequent Memory (hit vs. miss) | -0.04 | 0.10 | -0.4 | 0.69 |
| | Age × Accuracy | -0.28 | 0.16 | -1.68 | 0.10 |
| | Age × Subsequent Memory | -0.03 | 0.10 | -0.33 | 0.74 |
| | Accuracy × Subsequent Memory | -0.03 | 0.10 | -0.31 | 0.76 |
| | Age × Accuracy × Subsequent Memory | -0.04 | 0.09 | -0.45 | 0.66 |
| Theta Power | Age | 0.47 | 0.22 | 2.15 | 0.04* |
| | Accuracy | -0.24 | 0.20 | -1.17 | 0.25 |
| | Subsequent Memory (hit vs. miss) | -0.12 | 0.04 | -3.12 | $3.0\times10^{-3}$** |
| | Age × Accuracy | 0.13 | 0.20 | 0.62 | 0.54 |
| | Age × Subsequent Memory | 0.03 | 0.04 | 0.78 | 0.44 |
| | Accuracy × Subsequent Memory | -0.06 | 0.04 | -1.44 | 0.16 |
| | Age × Accuracy × Subsequent Memory | 0.05 | 0.04 | 1.20 | 0.24 |
| Theta Bandwidth | Age | -0.16 | 0.14 | -1.14 | 0.26 |
| | Accuracy | 0.04 | 0.13 | 0.27 | 0.79 |
| | Subsequent Memory (hit vs. miss) | 0.15 | 0.13 | 1.17 | 0.25 |
| | Age × Accuracy | 0.02 | 0.13 | 0.12 | 0.90 |
| | Age × Subsequent Memory | -0.08 | 0.14 | -0.58 | 0.57 |
| | Accuracy × Subsequent Memory | 0.09 | 0.13 | 0.71 | 0.48 |
| | Age × Accuracy × Subsequent Memory | 0.02 | 0.12 | 0.14 | 0.89 |

* $p < 0.05$, ** $p < 0.01$

**Table S7. LME model estimate for individual differences on anterior MTL theta parameters.**

| Parameters | Variables | Beta | Std. Error | t | P-value |
|---|---|---|---|---|---|
| Theta Frequency | Age | 0.11 | 0.16 | 0.64 | 0.52 |
| | Accuracy | 0.42 | 0.15 | 2.77 | $6.6\times10^{-3}$** |
| | Subsequent Memory (hit vs. miss) | -0.03 | 0.07 | -0.46 | 0.65 |
| | Age × Accuracy | -0.21 | 0.17 | -1.25 | 0.21 |
| | Age × Subsequent Memory | -0.02 | 0.07 | -0.24 | 0.81 |
| | Accuracy × Subsequent Memory | 0.02 | 0.07 | 0.32 | 0.75 |
| | Age × Accuracy × Subsequent Memory | 0.06 | 0.07 | 0.89 | 0.37 |
| Theta Power | Age | 0.31 | 0.17 | 1.88 | 0.06 |
| | Accuracy | 0.25 | 0.15 | 1.66 | 0.10 |
| | Subsequent Memory (hit vs. miss) | -0.11 | 0.04 | -3.00 | $3.3\times10^{-3}$** |
| | Age × Accuracy | 0.31 | 0.17 | 1.89 | 0.06 |
| | Age × Subsequent Memory | -0.003 | 0.04 | -0.08 | 0.94 |
| | Accuracy × Subsequent Memory | -0.07 | 0.04 | -1.90 | 0.06 |
| | Age × Accuracy × Subsequent Memory | 0.02 | 0.04 | 0.54 | 0.59 |
| Theta Bandwidth | Age | -0.09 | 0.12 | -0.74 | 0.46 |
| | Accuracy | -0.20 | 0.11 | -1.77 | 0.08 |
| | Subsequent Memory (hit vs. miss) | 0.13 | 0.09 | 1.32 | 0.19 |
| | Age × Accuracy | -0.01 | 0.12 | -0.07 | 0.94 |
| | Age × Subsequent Memory | -0.12 | 0.10 | -1.15 | 0.25 |
| | Accuracy × Subsequent Memory | 0.02 | 0.09 | 0.22 | 0.83 |
| | Age × Accuracy × Subsequent Memory | -0.07 | 0.10 | -0.75 | 0.45 |

** $p < 0.01$

**Table S8. Multiple regression model estimate for age and recognition accuracy effects on anterior MTL ERPAC subsequent memory effects.**

| Region | Referenced time | Variables | Beta | Std. Error | t | P-value |
|---|---|---|---|---|---|---|
| Anterior MTL | Response-related SME | Age | 0.42 | 0.25 | 1.66 | 0.12 |
| | | Accuracy | -0.23 | 0.24 | -0.96 | 0.35 |
| | | Age × Accuracy | 0.18 | 0.25 | 0.70 | 0.49 |

**Table S9. Multiple regression model estimate for age and recognition accuracy effects on posterior-anterior MTL theta phase coupling subsequent memory effects.**

| SME cluster | Variables | Beta | Std. Error | t | P-value |
|---|---|---|---|---|---|
| Scene-induced SME (Cluster 1) | Age | 0.41 | 0.25 | 1.61 | 0.14 |
| | Accuracy | 0.30 | 0.24 | 1.22 | 0.25 |
| | Age × Accuracy | 0.43 | 0.22 | 1.96 | 0.08 |
| Scene-induced SME (Cluster 2) | Age | 0.20 | 0.28 | 0.70 | 0.50 |
| | Accuracy | 0.37 | 0.28 | 1.35 | 0.21 |
| | Age × Accuracy | 0.28 | 0.25 | 1.11 | 0.29 |
| Response-related SME | Age | 0.49 | 0.23 | 2.13 | 0.06 |
| | Accuracy | 0.45 | 0.22 | 2.02 | 0.07 |
| | Age × Accuracy | 0.23 | 0.20 | 1.17 | 0.27 |

**Table S10. Correlation coefficients between posterior MTL HFA and anterior MTL theta parameters.**

| Posterior MTL HFA SME | Anterior MTL theta | Subsequent memory | r | p-value |
|---|---|---|---|---|
| Scene-induced | Theta frequency | Hit | 0.68 | $6.8 \times 10^{-3}$** |
| | | Miss | 0.65 | 0.01* |
| | Theta power | Hit | 0.30 | 0.30 |
| | | Miss | 0.05 | 0.86 |
| | Theta bandwidth | Hit | -0.20 | 0.50 |
| | | Miss | -0.44 | 0.11 |
| Response-related | Theta frequency | Hit | 0.60 | 0.02* |
| | | Miss | 0.36 | 0.21 |
| | Theta power | Hit | 0.37 | 0.19 |
| | | Miss | -0.13 | 0.67 |
| | Theta bandwidth | Hit | -0.27 | 0.35 |
| | | Miss | -0.07 | 0.80 |

* $p < 0.5$, ** $p < 0.01$
